# The sensor of the bacterial histidine kinase CpxA is a novel dimer of extracytoplasmic Per-ARNT-Sim (PAS) domains

**DOI:** 10.1101/2023.10.18.561931

**Authors:** Timothy H. S. Cho, Cameron Murray, Roxana Malpica, Rodrigo Margain-Quevedo, Gina L. Thede, Jun Lu, Ross A. Edwards, J. N. Mark Glover, Tracy L. Raivio

**Affiliations:** Departments of Biological Sciences; Biochemistry, University of Alberta, Edmonton, Alberta, Canada, T6G 2E9; Instituto de Ciencias Biomédicas, Universidad Autónoma de Ciudad Juárez, Chihuahua, México, 32310

**Keywords:** bacterial signal transduction, two-component systems, histidine kinase, envelope stress responses, AlphaFold2

## Abstract

Histidine kinases are key bacterial sensors that recognize diverse environmental stimuli. While mechanisms of phosphorylation and phosphotransfer by cytoplasmic kinase domains are relatively well-characterized, the ways in which extracytoplasmic sensor domains regulate activation remain mysterious. The Cpx envelope stress response is a conserved Gram-negative two-component system which is controlled by the sensor kinase CpxA. We report the structure of the *Escherichia coli* CpxA sensor domain (CpxA-SD) as a globular Per-ARNT-Sim (PAS)-like fold highly similar to that of *Vibrio parahaemolyticus* CpxA as determined by X-ray crystallography. Because sensor kinase dimerization is important for signaling, we used AlphaFold2 to model CpxA-SD in the context of its connected transmembrane domains, which yielded a novel dimer of PAS domains possessing a distinct dimer organization compared to previously characterized sensor domains. Gain of function *cpxA*\* alleles map to the dimer interface, and mutation of other residues in this region also leads to constitutive activation. CpxA activation can be suppressed by mutations that restore inter-monomer interactions, suggesting that inhibitory interactions between CpxA-SD monomers are the major point of control for CpxA activation and signaling. Searching through hundreds of structural homologues revealed the sensor domain of *Pseudomonas aeruginosa* sensor kinase PfeS as the only PAS structure in the same novel dimer orientation as CpxA, suggesting that our dimer orientation may be utilized by other extracytoplasmic PAS domains. Overall, our findings provide insight into the diversity of the organization of PAS sensory domains and how they regulate sensor kinase activation.

**Significance:** Bacterial two-component systems play an essential role in sensing environmental cues, mitigating stress, and regulating virulence. We approach the study of a key Gram-negative sensor kinase CpxA with both classical methods in structural biology and genetic analysis and emerging protein-folding prediction software. This approach provides a wholistic perspective on the structure and function of histidine kinases as proteins with modular and cellular compartment-spanning domain architectures. We report a novel organization of PAS domains in CpxA, highlighting the versatility and diversity of this sensory fold. Ultimately, these studies will facilitate the continued development of novel antimicrobials against sensor kinases, including CpxA, which is a previously studied target for antimicrobials.

## Introduction

Two-component systems (TCS) are ubiquitous bacterial sensory systems that recognize diverse environmental and cellular signals. Here, sensor histidine kinases sense stimuli and phosphorylate a cytoplasmic response regulator, which are usually transcription factors modulating the expression of target genes. Sensor kinases possess diverse and modular organizations^1–3^. Upon receiving a signal by sensory domains, conformational changes in these domains trigger downstream signaling events through common signal transduction domains, ultimately leading to autophosphorylation and phosphotransfer mediated by conserved kinase domains. While the structure and activity of cytoplasmic kinase domains are conserved and well-characterized, extracellular/periplasmic sensor domains and the mechanisms by which they sense and transduce signals are relatively more diverse^4,5^. Thus, the molecular basis for sensing and signal transduction by these domains remains more difficult to precisely characterize.

While extracytoplasmic sensor domains adopt a variety of folds, many sensor kinases (and other sensory proteins such as chemotaxis proteins) possess Per-ARNT-Sim (PAS)-like domains in this region^6^. Like other PAS domains, these extracytoplasmic PAS domains are characterized by a 5-stranded β-sheet with a 2-5-1-4-3 topology^7–9^. While often adopting highly similar folds, these domains tend to share low sequence homology^10^. Largely unique to extracytoplasmic PAS domains is the presence of a long N-terminal helix, sometimes termed the periplasmic helix (or p helix)^2^, which is continuous with the first transmembrane domain. These helices are thought to play a key role in transducing signals to downstream elements of sensor kinases (reviewed in ^2,11^), which largely exist and function as homodimers. Importantly, this helix forms the dimer interface in the prototypical periplasmic sensor domains of PhoQ^12^, DcuS^13^, and CitA^14^. Some authors have proposed that extracytoplasmic PAS-like domains be classed into a separate family based on these representative members called PhoQ-DcuS-CitA (PDC) domains due to common differences between intracellular and extracellular/periplasmic PAS domains^12^. However, the exact classification of these domains remains somewhat controversial. Recently, Upadhyay and colleagues proposed that extracytoplasmic PAS domains belong to a homologous but distinct family of domains known as Cache domains according to computational modelling^15^. Others have disputed whether extracytoplasmic PAS domains belong to a separate class of PAS domains because of the high level of sequence diversity between PAS domains and the high degree of structural similarity of the central β-sheets of intra- and extracellular PAS domains^8,16^. This diversity in nomenclature not only reflects a need for consensus in classification but also underscores the need for further study of these ubiquitous sensor domains.

CpxA is the sensor kinase of the CpxRA system, a conserved Gram-negative envelope stress response that responds to perturbations to envelope protein homeostasis^17,18^. Early studies of the Cpx response identified several alleles of *cpxA* (*cpxA\**) which possess constitutively activated phenotypes^19–21^. These *cpxA\** mutations were later sequenced and found to map to all regions of the protein^22^. Mutations from these studies mapping to the periplasmic portion of CpxA resulted in both hyper-activated and signal blind phenotypes, which led to the identification of this region as the main sensory domain. The periplasmic domain of CpxA in *Vibrio parahaemolyticus* was solved by X-ray crystallography as a globular PAS fold^23^. Like other extracytoplasmic PAS domains, the periplasmic domain of *V. parahaemolyticus’* CpxA contains a long α-helix at its N-terminus and a five-stranded β-sheet with the canonical 2-5-1-4-3 strand order. However, unlike most other sensor kinases, no known small molecule ligands have been found to bind CpxA. Instead, CpxA senses a wide variety of cues that result from envelope stress, especially those arising from misfolded proteins that affect inner membrane integrity^17,18,24,25^. CpxA also integrates signals from other envelope proteins such as the periplasmic chaperone-like protein CpxP^26–28^ and the outer membrane lipoprotein NlpE^29,30^. These factors help CpxA sense cues such as copper exposure and surface adhesion in the case of NlpE^30–33^ or the presence of misfolded pilus subunits in the case of CpxP^34–36^. However, NlpE and CpxP are not required for sensing many, if not most, Cpx inducing cues^37^, suggesting that they are not integral to CpxA’s inherent mechanism of sensing and activation.

While the structure of the *V. parahaemolyticus* CpxA periplasmic domain has been solved^23^, the structural and molecular basis for CpxA sensing and activation remains elusive. In addition, the Cpx response of *Vibrio* spp. remains relatively poorly characterized compared to studies of the system in *Escherichia coli* and related Enterobacteriaceae, and significant differences exist between signals sensed by CpxA in *Vibrio cholerae* and *E. coli*^38,39^. We report the structure of the *E. coli* CpxA periplasmic sensor domain (CpxA-SD_EC_) as determined by X-ray crystallography. CpxA-SD_EC_ adopts a globular PAS fold like the previously reported structure. To better understand CpxA-SD in its dimeric context, we used AlphaFold2 to model CpxA-SD. The resulting model predicted a novel organization of PAS domains in sensor kinases. Previously identified hyper-activated *cpxA\** alleles map to the modelled dimer interface, and mutation of conserved residues in this region also possess constitutively active and signal-blind phenotypes. The hyperactivation of these mutations can be completely or largely suppressed by introducing mutations predicted to restore interactions between CpxA-SD monomers, suggesting that CpxA kinase activity is controlled by inhibitory interactions between sensor domain monomers in the absence of inducing cues. Finally, the previously solved structure of PfeS (PDB 3KYZ), an enterobactin sensor from *Pseudomonas aeruginosa*, adopts highly similar characteristics to CpxA-SD. Taken together, we suggest that CpxA-SD represents a novel class of PAS domains that evolved to sense signals using a distinct dimer orientation from other PAS domains.

## Results

### CpxA-SD adopts a PAS fold

We purified and crystallized the *E. coli* CpxA sensor domain corresponding to residues 31-163 (CpxA-SD) and determined its structure to a final resolution of 1.8 Å (see supplemental methods and Table S1 for a full list of refinement statistics). Like CpxA in *Vibrio parahaemolyticus* (CpxA-SD_Vib_), *E. coli* CpxA-SD (CpxA-SD_EC_) adopts a PAS-like fold with each globular domain consisting of three α-helices surrounding a five-stranded antiparallel β-sheet in the canonical 2-5-1-4-3 topology (Figure 1A). Like other extracytoplasmic PAS domains, CpxA-SD, contains a long N-terminal helix (α1). However, this helix does not form the extreme N-terminus of the structure, instead containing an N-cap motif at its N-terminus and an extended tail region that folds against the β3 strand of the main PAS β-sheet (see later in Results for further discussion of this region). Overall, the monomer structure of CpxA-SD_EC_ is similar to that of the CpxA-SD_Vib_, with a root-mean-square deviation (RMSD) of 2.9 Å over 108 atoms (Figure 1B). As seen with other PAS domains^10^, CpxA-SD_EC_ and CpxA-SD_Vib_ secondary structure bears significant similarity despite a low sequence identity of 20% between residues 35-150. The regions around the α2 helix and the N-terminus of the α3 helix appear to be slightly different between these two structures. However, both regions appear to be affected by crystal packing, so the degree to which these differences are physiological is difficult to determine.

**Figure 1.**
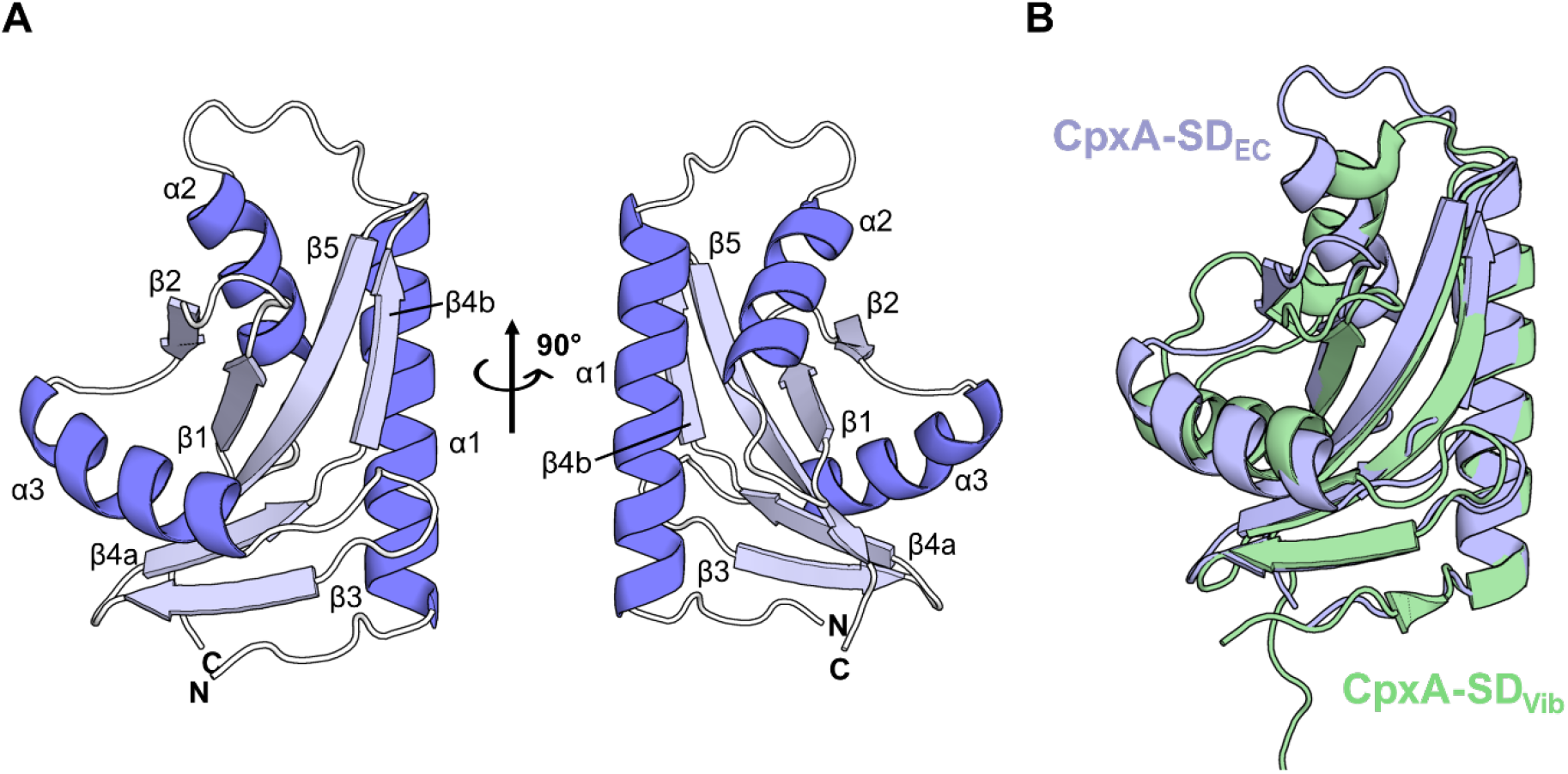
The sensor domain of CpxA adopts a PAS fold. **(A)** The resolved structure of CpxA-SD (aa 31-163) is shown as ribbon diagrams with secondary structure elements labelled α for helices and β for strands of β-sheets. Two views at a 90° rotation are shown. Shown is Chain B of the asymmetric unit. **(B)** Overlay of *E. coli* CpxA-SD (this study; blue) and CpxA_Vib_ (PDB 3V67; green).

### Conserved residues regulate basal CpxA activity and signal sensing

Several residues in the CpxA-SD appear to be conserved across hundreds of organisms. While most of these are hydrophobes buried within the PAS domain, asparagine 107 (N107), lysine 121 (K121), and tyrosine 123 (Y123) are notable exceptions (Figure 2A). These residues are also located within the deleted region of the constitutively activated and signal blind *cpxA24* allele (Δ92-123)^22^ (Figure 2B) and form a hydrogen bond network connecting the α3 helix and β3 strand in our crystal structure (Figure 2C). We hypothesized that this coordination may be important for CpxA activation. To investigate this, we introduced alanine swaps of these residues in the chromosomal copy of *cpxA* and tested the ability of these mutants to sense known Cpx activating signals, alkaline pH and NlpE overexpression (Figure 2D,E). Mutating these residues led to higher levels of basal activation of the Cpx response. The K121A and Y123A mutations also reduced sensitivity to alkaline pH and NlpE overexpression, reminiscent of previously identified *cpxA\** alleles^22^. CpxA N107A was still activated by these cues, albeit relatively more weakly compared to WT CpxA. While the protein levels of CpxA Y123A were similar to WT CpxA, CpxA N107A, and K121A showed slightly reduced expression levels (Figure S1). Overall, the results indicate that these residues are important for regulating CpxA activation, and by extension that interactions affecting α3 positioning play a key role here. Alternatively, the activation observed from CpxA N107A, K121A, and Y123A or the *cpxA24* deletion could be explained by misfolding of the sensor domain.

**Figure 2.**
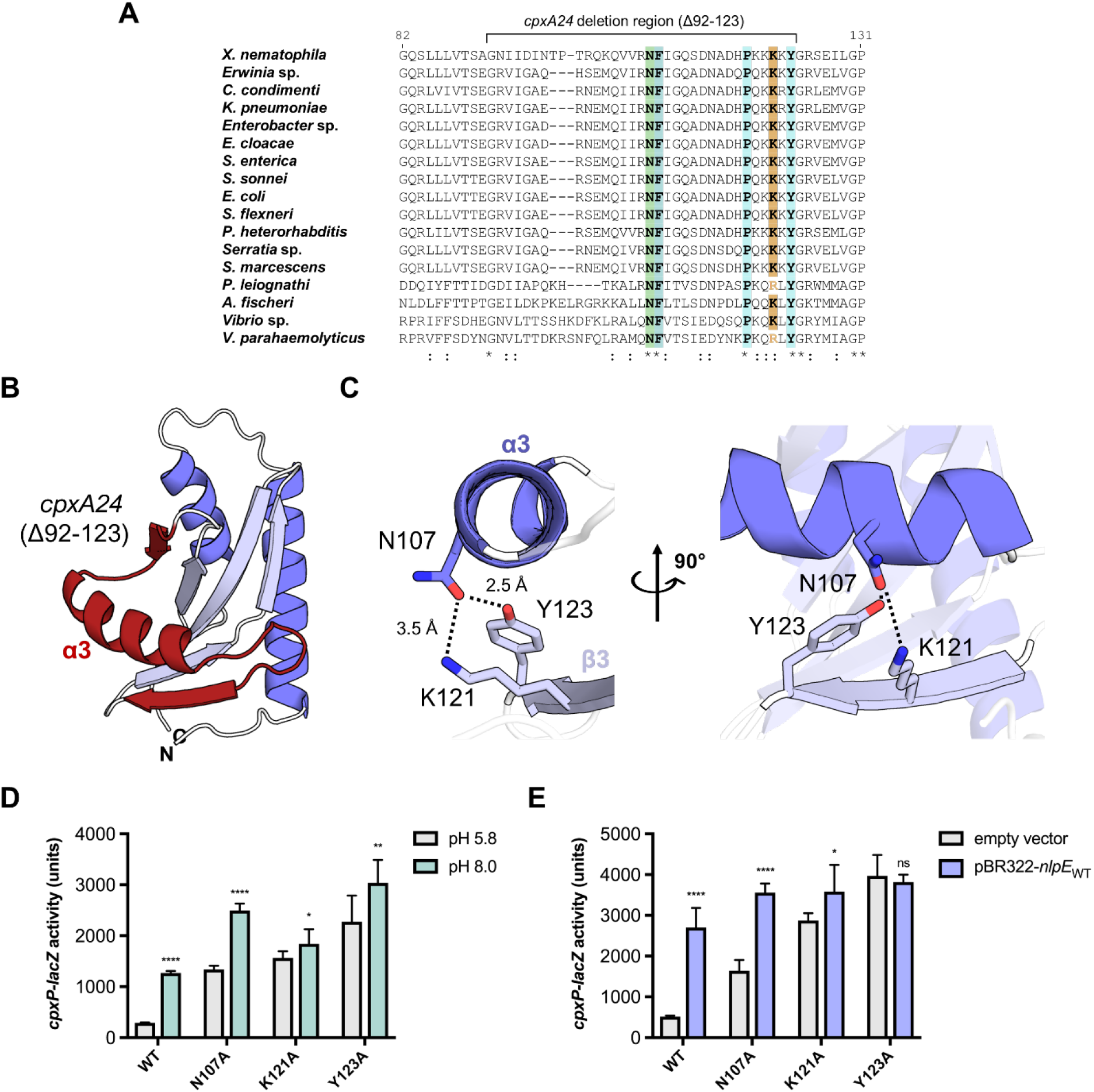
Conserved residues in CpxA-SD impact activation. **(A)** Multiple sequence alignment of CpxA-SD sequences from multiple organisms shows the conservation of several key residues in the *cpxA24* deletion region (Δ92-123), which is shown on the crystal structure of CpxA-SD in **(B)**. The hydrogen bond network formed by conserved residues N107, K121, and Y123 are shown in **(C)** in two views. Strains expressing chromosomal *cpxA* mutants (N107A, K121A, and Y123A) were tested for their ability to sense alkaline pH **(D)** and NlpE overexpression **(E)**, as measured by the activity of a chromosomal *cpxP-lacZ* reporter. Reporter activity was quantified by measuring β-galactosidase assay.

We wondered if the dimer structure of CpxA-SD would provide further insight into CpxA signaling as dimerization is important for sensor kinase structure and function^2,12,40^. The asymmetric unit of the crystal provided a dimer structure of CpxA-SD where the main dimer interface centres around its ɑ1 helices, which cross each other roughly perpendicularly (Figure S2A). This structure is somewhat reminiscent of an early structure of the *Salmonella enterica* PhoQ sensor domain dimer possessing similar crossed ɑ1 helices (PDB 1YAX)^41^, which appears to not be physiologically relevant^12,42^. In line with this, mutation of the M48 residue (M48K), which lies at the main dimer interface of this structure, did not lead to significant changes in activation of CpxA or its ability to sense alkaline pH (Figure S2B) or NlpE overexpression (Figure S2C), and CpxA M48K was expressed to similar levels as WT CpxA (Figure S2D). This was surprising given the importance of sensor kinase dimerization and led us to conclude that this dimer orientation was likely a crystallographic artifact.

### AlphaFold2 predicts a novel PAS domain dimer organization for CpxA-SD

While sensor kinases are extensively dimerized proteins, the presence of distinct domains in extracytoplasmic, membrane-integral, and cytoplasmic regions of the cell presents significant challenges to studying sensor kinases holistically. To investigate the dimeric structure of CpxA, we used the ColabFold implementation of AlphaFold2^43,44^ using custom multiple sequence alignments (MSA; see Methods) to model CpxA homodimers. Full length CpxA dimer models were initially generated (data not shown); however, monomers were rotationally symmetric and identical due to limitations in AlphaFold2. This artifactual symmetry resulted in poor alignments between the model and previously crystallized CpxA kinase domains, which possess significant asymmetry^45^ (data not shown). Additionally, any single model AlphaFold2 generates is an average of multiple potential states of CpxA (e.g. active and inactive)^43^. This averaging is exacerbated when the difference between the states is greatest, as is seen in the active and inactive states of cytoplasmic kinase domains^45^. To mitigate these issues, we excluded the histidine phosphotransfer (DHp) and catalytic (CA) domains from our modeled complexes, limiting our models to the region encompassing transmembrane helix 1 (TM1), the PAS domain, transmembrane helix 2 (TM2), and the HAMP domain (Figure 3A). These smaller models had higher local (pLDDT) and global (pTm) confidence than the full length models, which indicated that these models better represent a single state of CpxA (Figure S3)^43^. Both *E. coli* and *V. parahaemolyticus* CpxA models were created either terminating at the end of TM2 or the HAMP domain. The top dimer structures were similar for both the *E. coli* and *V. parahaemolyticus* structures, and the inclusion of the HAMP domain did not significantly influence the overall dimer arrangement (Figure 3B,C, S4). Notably, the sensor PAS domain modelled by AlphaFold2 was virtually identical to the structure we solved by X-ray crystallography (Figure S5).

**Figure 3.**
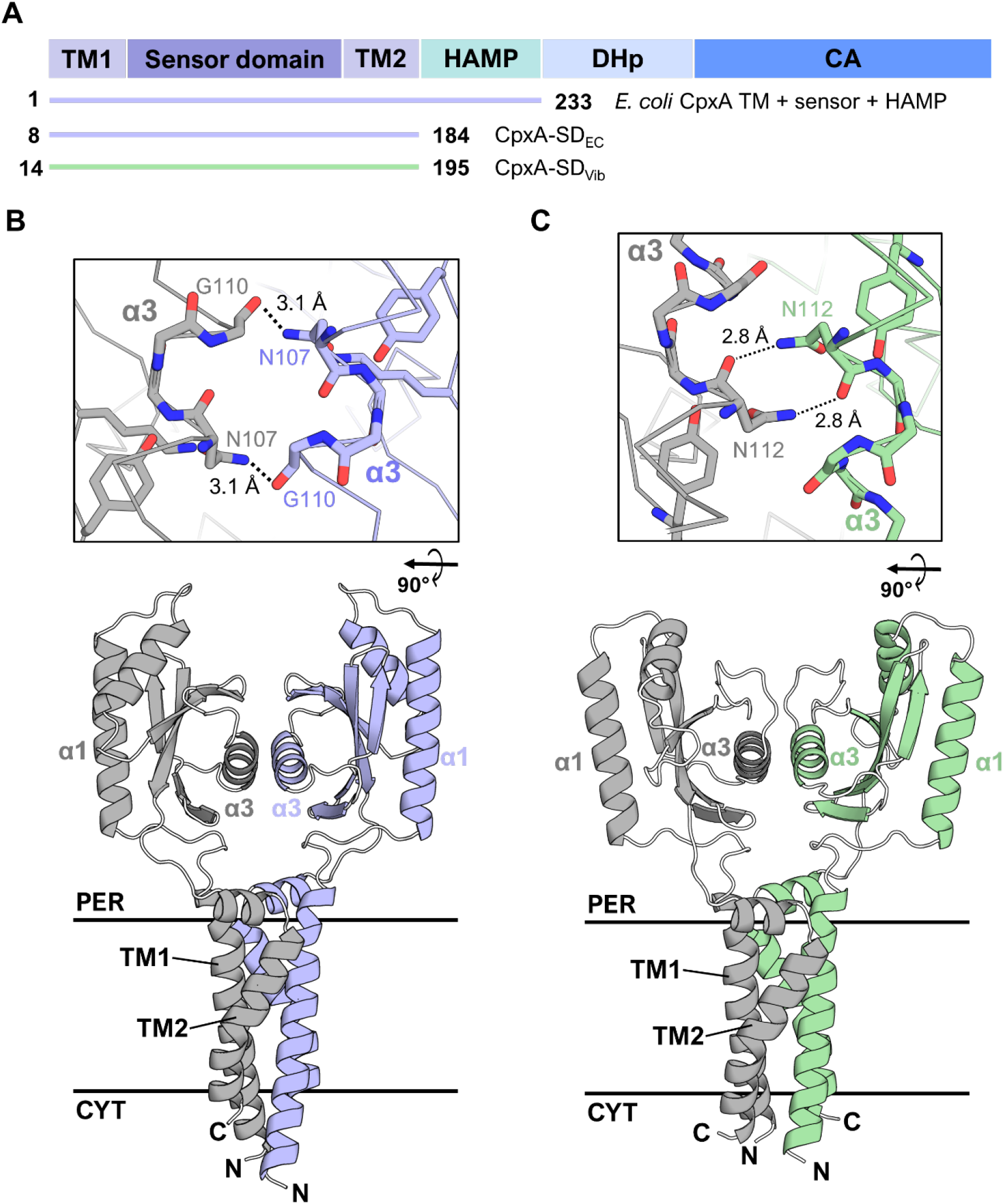
AlphaFold2 models of CpxA sensor and transmembrane domain dimers. **(A)** Domain architecture of CpxA with the AlphaFold2-modeled portions of each CpxA sequence from *E. coli* (*E. coli* CpxA TM + sensor + HAMP, CpxA-SD_EC_) and *V. parahaemolyticus* (CpxA-SD_Vib_). Modelled amino acids numbers are shown alongside each schematic of the modeled region of CpxA. AlphaFold2 models of the sensor and transmembrane (TM1 and TM2) domains of **(B)** *E. coli* and **(C)** *V. parahaemolyticus* CpxA. The α3-α3 dimer interface of both proteins (from a top-down view) is shown in the insets above each of the structures.

Both CpxA-SD_EC_ and CpxA-SD_Vib_ were modeled as homodimers with their TM domains in a tight four helix bundle, similar to other histidine kinases (Figure 3B,C)^46,47^. However, their PAS domains were oriented opposite to those found in most histidine kinases, which are oriented such that the α1 helices form the dimer interface, while the α3 helices are exposed and (usually) involved in ligand binding^9^. In contrast, the CpxA-SD PAS domains were modelled with the α3 helices forming the dimer interface, placing N107, K121, and Y123 at the interacting surface. The aforementioned *cpxA24* mutation encompasses nearly the entirety of the α3 helix and β3 strand, which comprises the majority of the CpxA-SD dimer interface in the AlphaFold2 model. Importantly, the previously identified N107 residue lies at the dimer interface in the AlphaFold2 models. N107, K121, and Y123 are structurally conserved in both *E. coli* and *V. parahaemolyticus.* K121 and Y123 form a hydrogen bond network with N107 (Figure 2C), which has the effect of precisely positioning N107 such that its amine group protrudes towards the other monomer (top inset of Figure 3B). In both *E. coli* and *V. parahaemolyticus* models, this central asparagine (N107 and N112, respectively) forms hydrogen bonds with a main chain carbonyl group of the other monomer (top insets of Figure 3B,C). While the exact positioning of these hydrogen bonds is modeled differently between CpxA-SD in *E. coli* and *V. parahaemolyticus*, these bounds could limit the inter-monomer flexibility of this dimer orientation in both models. Thus, we reasoned that disrupting these interactions would destabilize the CpxA-SD dimer and impact CpxA activation.

We further tested the roles of the conserved N107, K121 and Y123 residues and their predicted interactions in *E. coli* CpxA. To facilitate further testing of *cpxA* mutants, we cloned *cpxA* into plasmid pK184 and introduced it into a strain lacking chromosomal *cpxA* and possessing a chromosomal CpxA-regulated *cpxP-lacZ* reporter (TR50). We confirmed that this plasmid-based system complements the phenotype of the Δ*cpxA* mutant as it restored the ability of the mutant to sense alkaline pH and NlpE overexpression (compare the WT reporter strain TR50 vs the Δ*cpxA* strain expressing CpxA [WT] in Figure 4). This was despite the fact that expression from the plasmid was significantly higher than native levels of CpxA (Figure S1). Using this system, we again looked at the function of N107. The higher activation of CpxA N107A suggests the AlphaFold2-predicted dimer interface was physiologically relevant. However, its phenotype could be explained by the alanine substitution disrupting the intra-monomer hydrogen bonding network (Figure 2C). Based on the model (Figure 3B), we predicted that replacing N107 with an aspartate (N107D) would yield a phenotype specific to interactions between CpxA-SD by introducing repulsion at the interface between monomers while maintaining the hydrogen bonding network between α3 and β2 within the monomer (Figure 2C). We also introduced mutations into neighbouring residues Gln103 and Arg106 which reside on the α3 helix and face towards the other CpxA-SD monomer in the AlphaFold2 model (Figure 4A). We found that CpxA N107D was hyper-activated in the absence of inducing cues and no longer sensed alkaline pH and NlpE overexpression, resulting in a signal blind phenotype (Figure 4B,C). Similarly, CpxA Q103E and R106E showed high levels of activation in the absence of inducing signal compared to WT CpxA and were insensitive to inducing cues (Figure 4B,C).

**Figure 4.**
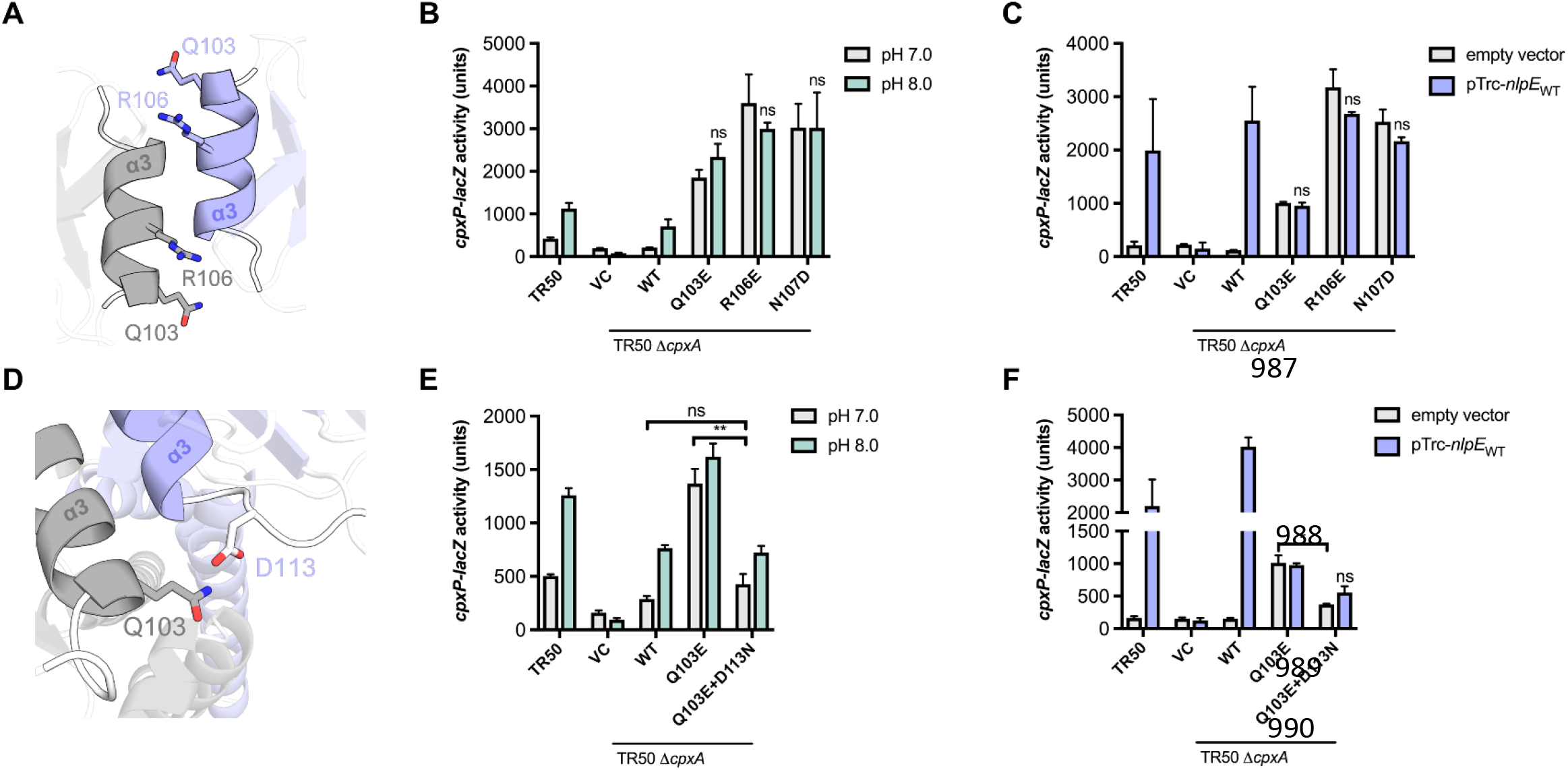
Mutations in the dimer interface of CpxA-SD lead to activation. **(A)** Other residues on the α3 helix. Residues Q103 and R106 are shown on the AlphaFold2 model of *E. coli* CpxA-SD. The ability of mutants of CpxA Q103, R106, and N107 in *E. coli* to sense alkaline pH and NlpE overexpression are shown in **(B)** and **(C),** respectively. TR50 refers to the WT strain of *E. coli* MC4100 encoding a chromosomal *cpxP-lacZ* reporter. Variants of CpxA were expressed from plasmids in a TR50 Δ*cpxA* background and tested for activation of a *cpxP-lacZ* reporter. **(D)** The position of Q103 and D113 on opposing monomers are shown. The ability of the rescued CpxA Q103E+D113K mutant to sense these signals are shown in **(E)** and **(F)**. Shown are a mean of three independent replicates. Significance indicates t-tests that were conducted between the bars shown in brackets or, in the case that no brackets are shown, compared to either pH 7.0 or empty vector treatments of the same genetic background (* p<0.05, ** p<0.01).

### Charge swap mutations at the predicted dimer interface restore signaling

Interestingly, the Q103E mutation appears to lock CpxA into a particular level of activation as this variant was not activated to the same extent basally as the other mutations, and NlpE overexpression did not activate CpxA Q103E to the levels seen either in WT CpxA or the other mutants. AlphaFold2 modelling predicted that Q103 could hydrogen bond to D113 on the opposing CpxA-SD monomer (Figure 4D), leading us to hypothesize that the Q103E mutation results in a charge repulsion between monomers that leads to aberrant CpxA activation. We tested if the hyper-activation of this mutant could be suppressed by re-introducing a hydrogen bond at this position via an asparagine substitution at D113. Strikingly, we observed that Q103E and D113N mutations together suppress the hyper-activation of CpxA Q103E in the absence of inducing cues (pH 7.0 or empty vector in Figure 4E,F). Further, CpxA Q103E+D113N was activated to a similar extent as WT in the presence of alkaline pH (pH 8.0). Interestingly, CpxA Q103E+D113N was largely insensitive to NlpE overexpression, similar to the CpxA Q103E single mutant. The inability of CpxA Q103E+D113N to sense NlpE overexpression may indicate that this interface is important for NlpE activation of CpxA, but a mechanism for this requires further investigation. As expected, introducing a D113K mutation into CpxA also confers a hyper-activated and signal blind phenotype (Figure S6), supporting our hypothesis that disrupting interactions between CpxA-SD monomers leads to inappropriate CpxA activation.

To further test the role of the predicted dimer interface in signaling, we used the AlphaFold2 model to investigate previously identified *cpxA\** alleles which map to the sensor domain. *cpxA102* is a single amino acid substitution of a lysine at E91^22^, which maps to the β2 strand that is located towards the top of the dimer interface in the AlphaFold2 model. Further examination of the residues surrounding this allele revealed several nearby positively charged arginines (R93 and R99) at the dimerization interface (Figure 5A). While none of these residues are predicted to form hydrogen bonds with E91, we hypothesized that the introduction of a positive charge in this region may introduce repulsion between CpxA-SD monomers, leading to the observed constitutive activation. Based on this hypothesis, we attempted to rescue the hyper-activation of CpxA E91K by introducing negatively charged residues at R93 and/or R99.

**Figure 5.**
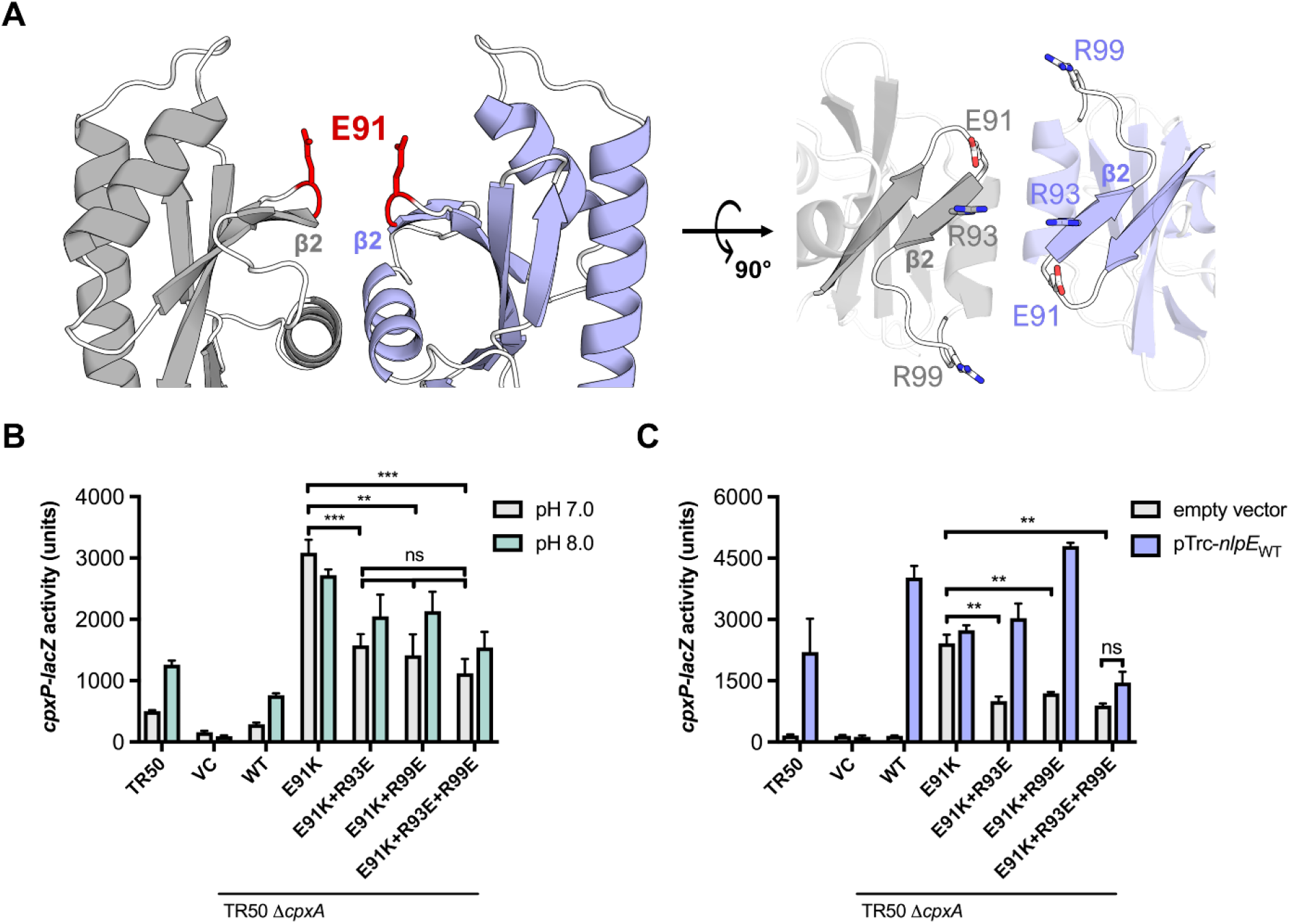
Other mutations of CpxA-SD lead to hyper-activation. **(A)** shows the location of E91, the residue mutated in the *cpxA102* allele (CpxA E91K). Two views are shown, with surrounding basic residues R93 and R99 shown in the right panel. The ability of E91K-containing mutations expressed from plasmid in a Δ*cpxA* strain to sense **(B)** alkaline pH and **(C)** NlpE overexpression was tested in by assaying the activity of a *cpxP-lacZ* reporter. Shown are a mean of three independent replicates (t-test * p<0.05, ** p<0.01, *** p<0.001).

We introduced the E91K mutation singly as well as in combination with mutations predicted to restore wild-type charge to the dimer interface (R93E and R99E) into our CpxA expression vector and measured both the basal activity of these variants and their ability to respond to alkaline pH and NlpE overexpression (Figure 5B,C). Consistent with previous studies^22^, we found that CpxA E91K was hyper-activated basally at pH 7 (Figure 5B) and in the absence of NlpE overexpression (Figure 5C). CpxA E91K was also insensitive to alkaline pH and NlpE overexpression (Figure 5B,C). In line with our hypothesis, the introduction of either R93E or R99E in combination with E91K into CpxA was significantly reduced basal activation, although not completely to wildtype levels. Interestingly both mutations reduced activation to similar extents and there was no additive effect of introducing these mutations in tandem, as the triple mutant (CpxA E91K+R93E+R99E) possessed similar levels of basal activation to the double mutations. These rescued mutations slightly restored the ability of CpxA to sense alkaline pH, as shown by slightly higher activity at pH 8; however, they were significantly less sensitive to pH than WT CpxA. CpxA E91K+R93E and E91K+R99E were sensitive to NlpE overexpression. In contrast, the triple mutant was largely insensitive to it.

Unlike Q103 and D113, these residues are near the previously reported NlpE binding interface^48^ (Cho et al. in preparation) so they could interfere with the binding of NlpE to CpxA; however, this requires further investigation. CpxA expression levels are unlikely to account for any of the observed phenotypes since all of these mutants are expressed at similar levels to WT CpxA from our plasmid (Figure S1). Further, introducing an E91A mutation into CpxA, while leading to higher activation, did not lead to the same level of hyper-activation and signal-blindness as the CpxA E91K charge swap (Figure S7). Thus, constitutive activation of *cpxA102* (CpxA E91K) is likely caused by the repulsion introduced by the lysine residue with neighbouring positive residues between CpxA-SD monomers.

### CpxA-SD is a novel dimer of extracytoplasmic PAS domains

These findings suggest that, like the AlphaFold2 model, the α3 helix of the CpxA-SD PAS domain is proximal to the dimer interface while CpxA’s α1 helix is distal to the dimer interface. As mentioned previously, this dimer orientation is unusual and, to our knowledge, has not been reported. We verified this by performing a structural homology search using the web server Dali^49^ and the crystal structure of CpxA-SD as a query. This search yielded 700 non-unique molecules with a Z-score greater than two. Among the top hits were the prototypical extracytoplasmic PAS domains of the sensor kinases PhoQ, CitA, and DcuS (Figure 6A-C), and other PAS domain sensors (Figure S8). While these structures also feature long α1 helices, their α1 helices are at the dimer interface and are likely continuous with their respective transmembrane domains^6^. A custom python script was used to search the homologous structures to find any that exhibit a similar dimer orientation to that found in the CpxA AlphaFold2 model. For each PDB entry, each pair of nearby chains in that entry were evaluated to find parallel dimers where the α3 helices are facing each other.

**Figure 6.**
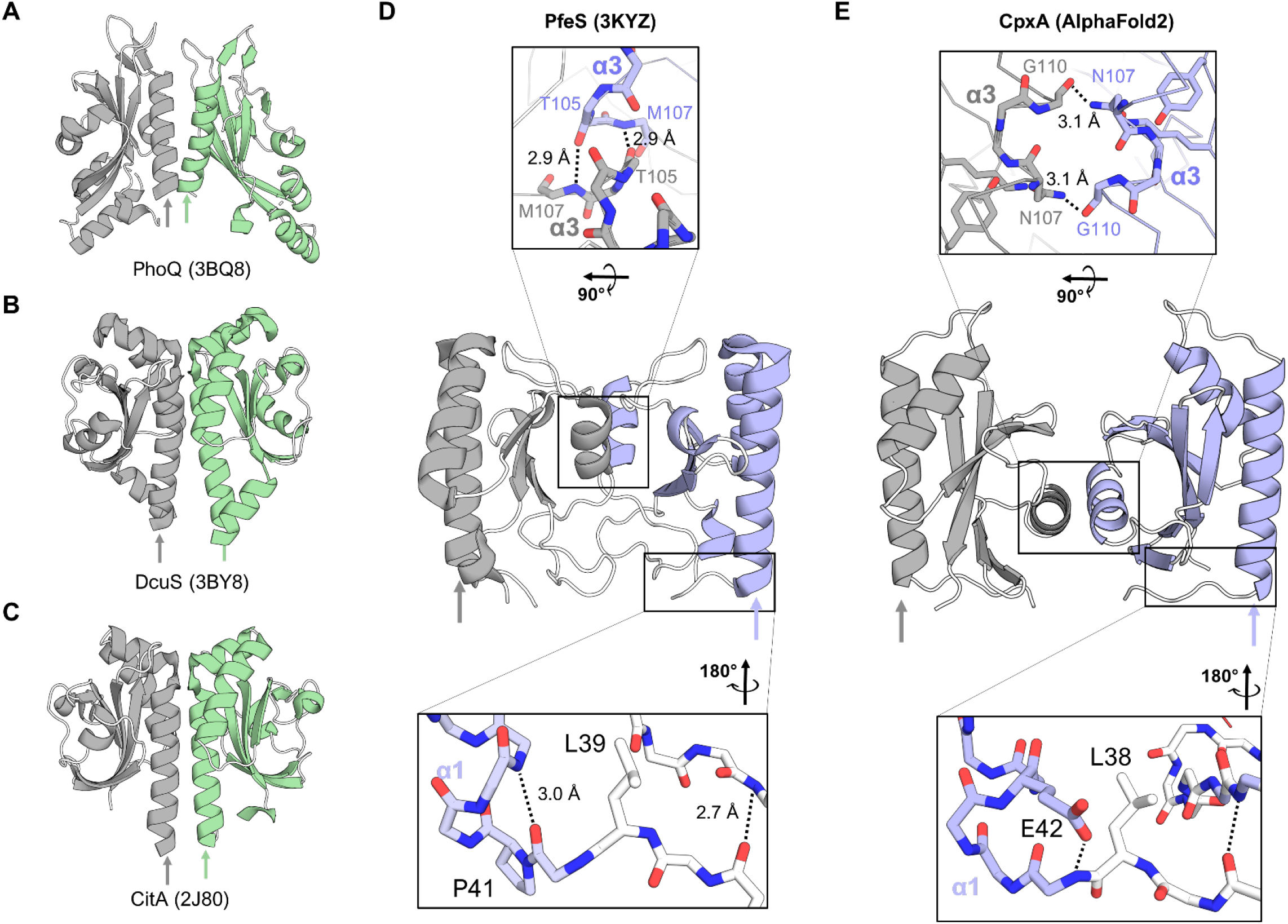
The PAS domains of CpxA adopt a novel orientation. The structures of the representative PAS domains of histidine kinases: **(A)** *E. coli* PhoQ (PDB 3BQ8), **(B)** *Klebsiella pneumoniae* CitA (PDB 2J80), **(C)** *E. coli* DcuS (PDB 3BY8). The crystal structure of the periplasmic domain of enterobactin sensor PfeS (PDB 3KYZ) from *Pseudomonas aeruginosa* is shown in **(D)** with key areas shown as insets above (dimer interface) and below (“bridge”) the structure. A parallel comparison to the AlphaFold2 model of *E. coli* CpxA-SD is shown in **(E)**, with corresponding regions also shown in insets.

One hit from our search had striking similarity to CpxA-SD_EC_: the crystal structure of the periplasmic sensor domain of PfeS (PDB 3KYZ), a sensor of enterobactin from *Pseudomonas aeruginosa* (Figure 6D)^50,51^. The PfeS dimer interface also centres on its α3 helix while its α1 helix is oriented towards the outside of the protein. At the dimer interface, symmetrical hydrogen bonds between the α3 helices (T105-M107 in the case of PfeS, N107 in CpxA; top insets of Figure 6D and E) are present near the axis of rotation, and the N-terminus of the PfeS α1 helix possesses a proline containing N-cap preceded by a leucine in the same orientation as CpxA-SD_Vib_ and CpxA-SD_EC_ (compare bottom insets of Figure 6D and E). Since an N-cap at the α1 helix appeared to be characteristic of this dimer form, a second search of the Dali hits was performed to find monomers that possessed such an N-cap. This search revealed many α1 N-cap containing PAS structures, including soluble and membrane-bound histidine kinases. While most of the histidine kinases were crystallized as α1-helix dimers, one structure of the complex carbohydrate sensor AbfS from *Cellvibrio japonicus* (PDB 2VA0)^52^ was not and represents another potential α3-helix-dimer candidate, suggesting novel CpxA-like PAS dimers may be more prevalent across histidine kinases.

We propose that the extracytoplasmic PAS domains of CpxA, while adopting essentially the same overall fold as other PAS domains, has evolved to adopt a distinct dimer conformation which may be present in other PAS domain-containing histidine kinases. Interestingly, CpxA, like PfeS, has been implicated in sensing enterobactin or enterobactin-related signals; periplasmic accumulation of enterobactin or vibriobactin is thought to strip iron from membrane-integral iron-sulfur cluster-containing proteins, disrupting protein folding and activating CpxA^53,54^. While PfeS appears to sense enterobactin to combat iron starvation^50,51^, the structural and sensory parallels between these proteins may point to a signaling mechanism unique to this dimeric arrangement of PAS domains.

## Discussion

The extracytoplasmic PAS domains of histidine kinases play a key role in signal sensing and controlling kinase activity. In this study, we report the crystal structure of CpxA’s PAS domain and use AlphaFold2 to investigate how this domain regulates CpxA activation. Our results suggest that interactions between residues at the novel CpxA-SD PAS dimer interface regulate CpxA’s kinase activity. Because disturbance of this interface almost invariably results in aberrant CpxA activation, the basal state of the CpxA kinase domains is likely kept OFF by inhibitory interactions between CpxA-SD monomers. This is reminiscent of findings in other sensor kinases and sensory proteins. Lee and colleagues propose that the N-terminal PAS (PAS-A) domain of the sporulation regulator KinA of *Bacillus subtilis* regulates kinase activation by its dimerization state^55^. In a model with many parallels to our own, dimeric PAS-A represents a basal or kinase off state of KinA, while monomeric PAS-A corresponds to activation of KinA. This model was based on studies of bacterial phytochromes which found that in activated proteins, N-terminal sensor domains are distal, while being dimeric in inactive conformations^56^. Despite the fact that both of these proteins are cytoplasmic, soluble histidine kinases, their extensive dimerization is shared with membrane-bound histidine kinases. While there appears to be some controversy about the exact contribution of the PAS-A domain to KinA function^57^, this model nonetheless provides an interesting parallel to our own model of how PAS sensor domain dimerization regulates activation in the context of a highly dimerized protein.

Previous studies have suggested that CpxA is biased towards an OFF state by interaction with the periplasmic chaperone-like protein CpxP. Evidence supports a model where CpxA is shifted towards the active state by sequestration of CpxP through its interaction with misfolded protein substrates^28^. However, deletion of *cpxP* does not lead to full activation of the Cpx response, and inhibition of CpxA by CpxP overexpression does not occur in *cpxA\** strains^27^. In line with these results, we found that hyper activating mutations in the sensor domain, such as CpxA E91K and D113K, are insensitive to CpxP overexpression (Figure S9). Further, ratio of CpxA to CpxP present in the periplasm is unlikely to be one to one. While CpxP and CpxA appear to be translated at similar levels^58^, the protein levels of CpxP are heavily regulated by proteolysis such that CpxP is only detectable by Western blot when DegP, the protease responsible for CpxP degradation, is deleted or if the Cpx response is strongly activated^35^. Because it is unlikely that there is a CpxP molecule for every CpxA molecule present in the envelope, it is unlikely that CpxP is fully responsible for maintaining CpxA in an OFF state, suggesting additional, CpxA-inherent mechanisms control the activation of CpxA in the absence of inducing cues.

The behaviour of our dimer interface mutants also warrants a contrasting comparison to the sensor domain of PhoQ. Dimerization of the sensor domain of PhoQ has been extensively characterized^12,42^; monomer-monomer interactions in the sensor domain of PhoQ appear to control PhoQ activation, as disrupting the dimer interface abrogates low magnesium sensing. However, it does not lead to constitutive activation like mutating CpxA’s dimer interface^12^. Thus, the dimer interface of PhoQ may play an important role in facilitating activation. In line with this, Mensa and colleagues observed that the PhoQ sensor domain appears to favour an “ON” state, as seen in experiments mutationally decoupling sensor, HAMP, and catalytic domains^59^. Because the dimer interface of CpxA-SD is sensitive to mutations, which tend to have activating effects, relatively modest dimer interface of CpxA-SD may increase sensitivity to signals present in the periplasm, such as the presence of a broad range of misfolded proteins or small changes in the levels of interacting partner proteins such as CpxP or NlpE. This sensitivity may also be linked to sensing deformations in the membrane such as defects in translocation through the Sec pathway^60^, respiratory complex assembly^61,62^, or changes in membrane composition^63,64^. However, the molecular basis for how CpxA senses misfolded proteins in the periplasm remains unclear. In comparison to other PAS domain containing sensor kinases, studies of CpxA may be hampered by the lack of known small molecule ligands for CpxA. Thus, future studies should focus on CpxA’s known protein partners, as direct binding of NlpE to CpxA appears to activate CpxA^31,48,65^. CpxP overexpression and interaction inhibits basal kinase activity of CpxA^22,27,66^, suggesting that CpxP may act predominantly as a factor to stabilize CpxA sensor domains to dull activation of CpxA in a negative feedback mechanism. Further structural studies of the sensor domain of CpxA in these contexts may shed more light on how the non-typical PAS orientation of CpxA regulates kinase activation.

Our work shows that previously characterized and structure-guided *cpxA*\* mutations are likely to alter signaling by impacting folding within monomers (Figure 2) and/or those between monomers at a novel dimer interface (Figures 3, 4). We therefore further investigated how other previously isolated *cpxA*\* mutations might affect the structure of CpxA, potentially providing insight into how CpxA transmits signals from the periplasm. Interestingly, we identified two domain-domain interface regions where clusters of *cpxA\** alleles have been found to map, namely, the PAS-TM interface and the TM-HAMP interface.

The crystal structures of CpxA-SD_EC_, CpxA-SD_Vib_, and PfeS all possess an α1 helix that is preceded by a seemingly unstructured tail (Figure 3B,C, 6). The existence of this region does not seem to be a coincidence since such a region would be required to accommodate the novel dimer orientation of CpxA while maintaining both the overall domain organization and tightly packed TM bundle exhibited in histidine kinases^2^. In the AlphaFold2 models, we see exactly that; this region links α1 to TM1 and forms the PAS-TM interface. The most notable similarity in the three crystal structures was an N-cap motif at the N-termini of the α1 helix. N-cap motifs are diverse in sequence and stabilize exposed N-terminal main chain amines of the helix (Figure 6D,E lower inset)^67^. These motifs are seen in 92.2% (296/321) of analyzed CpxA homologues, with the other 7.8% (25/321) possessing motifs that are found in known N-caps but are not traditional capping box motifs (Figure S10). The N-cap motif in all three structures is preceded by a leucine or isoleucine, which is present in 89.4% (287/321) of analyzed CpxA homologues with other branched hydrophobes (methionine and valine) making up a further 8.4% (27/321). The level of structural and sequence conservation suggests this is a critical region for protein function. Indeed, an L38P (*cpxA9*) mutation was reported to strongly activate CpxA kinase activity^19,68^, despite prolines being common in *cpxA* sequences at flanking residues 37 and 39. Based on structures of N-caps with proline at this position, it’s likely that this mutation promotes backbone conformers other than those seen in the crystal structure^67^, possibly decoupling the main PAS fold of CpxA from the first TM domain. Other small secondary structure elements exist N-terminal to the N-cap that bridge the PAS and TM1 domains in the AlphaFold2 models (Figure 7A). In CpxA-SD_EC_, these elements are tethered to TM2 via Q153 and the salt bridge between R33 and D162, which appears to be highly conserved (Figure 7A). Two other *cpxA\** alleles^22^, *cpxA104* (CpxA R33C) and *cpxA103* (CpxA R163P) likely act by disrupting this salt bridge either directly, in the case of *cpxA104* (CpxA R33C), or indirectly by significantly altering the position of D162 (CpxA R163P, *cpxA103*)^20,22^. Similar to our observations of the CpxA PAS domain dimer, our model suggests that interactions in this bridge region stabilize an OFF state. The fact that all of these mutations lead to aberrant activation of CpxA suggests that activation of CpxA likely involves signals being transduced from the PAS domain through this linker either directly to TM1 or indirectly to TM2.

**Figure 7.**
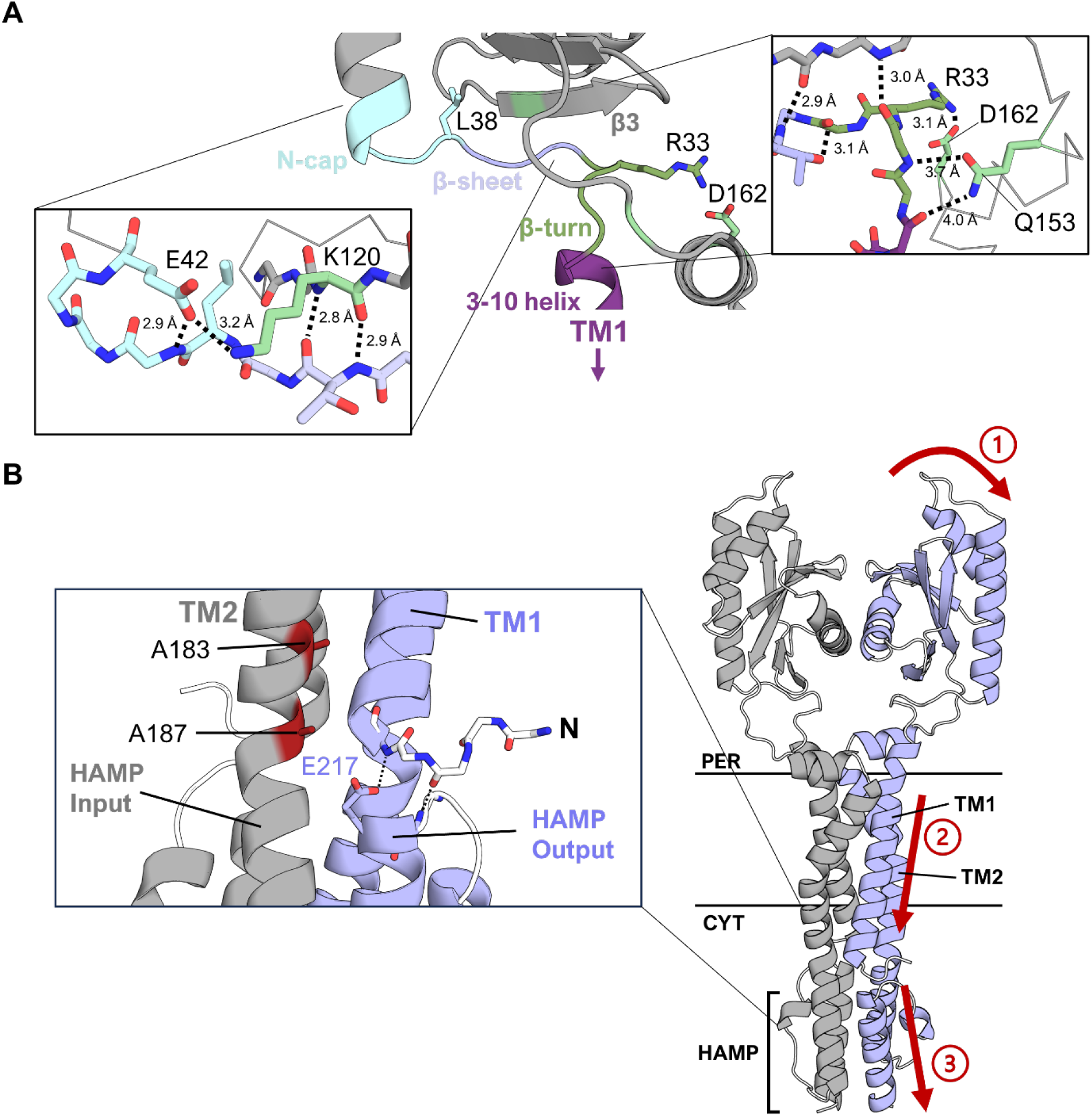
The AlphaFold2 model explains the phenotypes of mutants in other regions of CpxA. **(A)** The bridge region from the AlphaFold2 model of CpxA-SD_EC_ is shown with key elements colour coded. Insets show rotated and zoomed in stick models of those regions with pertinent predicted interactions shown with distance measurements. A 3-10 helix is a helix where the i-th residue hydrogen bonds to the i+3 residue instead of α helical i to i+4 hydrogen bonding. **(B)** The AlphaFold2 model of CpxA including the HAMP domain. Inset shows previously identified mutations neighbouring the HAMP domain which lead to activation of CpxA. Red arrows show a tentative mechanism of how signals are transduced from the sensor domain to the HAMP domains based on data presented in this paper as well as historical data.

Further insights into signal transduction by CpxA can be gleaned from mapping other *cpxA\** mutations onto the AlphaFold2 model including the cytoplasmic HAMP domains directly following the TM domains (Figure 7B). In this model, the N-terminus of the first TM domain packs against the top of the output helix of the HAMP domain. At the cytosolic end of the TM bundle, A183 and A187 are modeled to pack against TM1 of the opposing dimer close to where TM1 contacts the output helix of the HAMP domain (Figure 7B). These positions show strong selection for small residues such as Ala, Gly, Ser and Thr, resembling GxxxG motifs^69^. Previous studies have identified activating *cpxA\** mutations *cpxA711* (CpxA A183T) and *cpxA17* (CpxA A187E) in this region^20,22^. Disruption of this tight packing through mutations of A183T or A187E would perturb the modeled interaction between the highly conserved glutamate (E217) on the output helix of the HAMP domain and the N-cap of TM1. Breaking this conserved interface between signal transduction domains is likely to affect CpxA’s activity and would represent a large change from the AlphaFold2 model, which appears to represent a basal state of CpxA, implying that TM1 mobility is important for signal transduction from the periplasmic PAS domain. The AlphaFold2 models of CpxA_EC_ along with our, and previously published, mutation data suggest an overall signal path for CpxA. Disruption of interactions between the PAS domains causes signal to be transmitted through the PAS-TM1 linker to TM1, which carries the signal through the membrane. The N-terminus of TM1 interacts with the output helix of the HAMP domain which passes signal to the kinase domains (Figure 7B). This model will be a useful generalization to guide further study of signal transduction in CpxA and other histidine kinases.

## Conclusion

Despite the crucial role of sensor domains of histidine kinases in initiating kinase activation, considerable mystery remains as to how these sensor domains function due to both the diversity of the signals sensed by these proteins and the structures of their sensor domains^4,5^. While CpxA contains a PAS domain that is largely similar to the PAS folds of other sensor kinases, CpxA’s “non-canonical” dimer structure regulates its ability to sense periplasmic triggers (Figure 8); diverse inputs that disrupt the stability of the CpxA-SD dimer appear to activate CpxA kinase activity. This is clearly indicated by mutations that disrupt dimerization leading to activation, whereas mutations restoring dimer stability rescue CpxA’s basal OFF state. We hypothesize that activating signals might work by a similar mechanism of destabilizing the CpxA-SD dimer. Further work is required to test this model in the context of known CpxA inducing signals, such as the presence of unfolded proteins, high pH, and NlpE binding.

**Figure 8.**
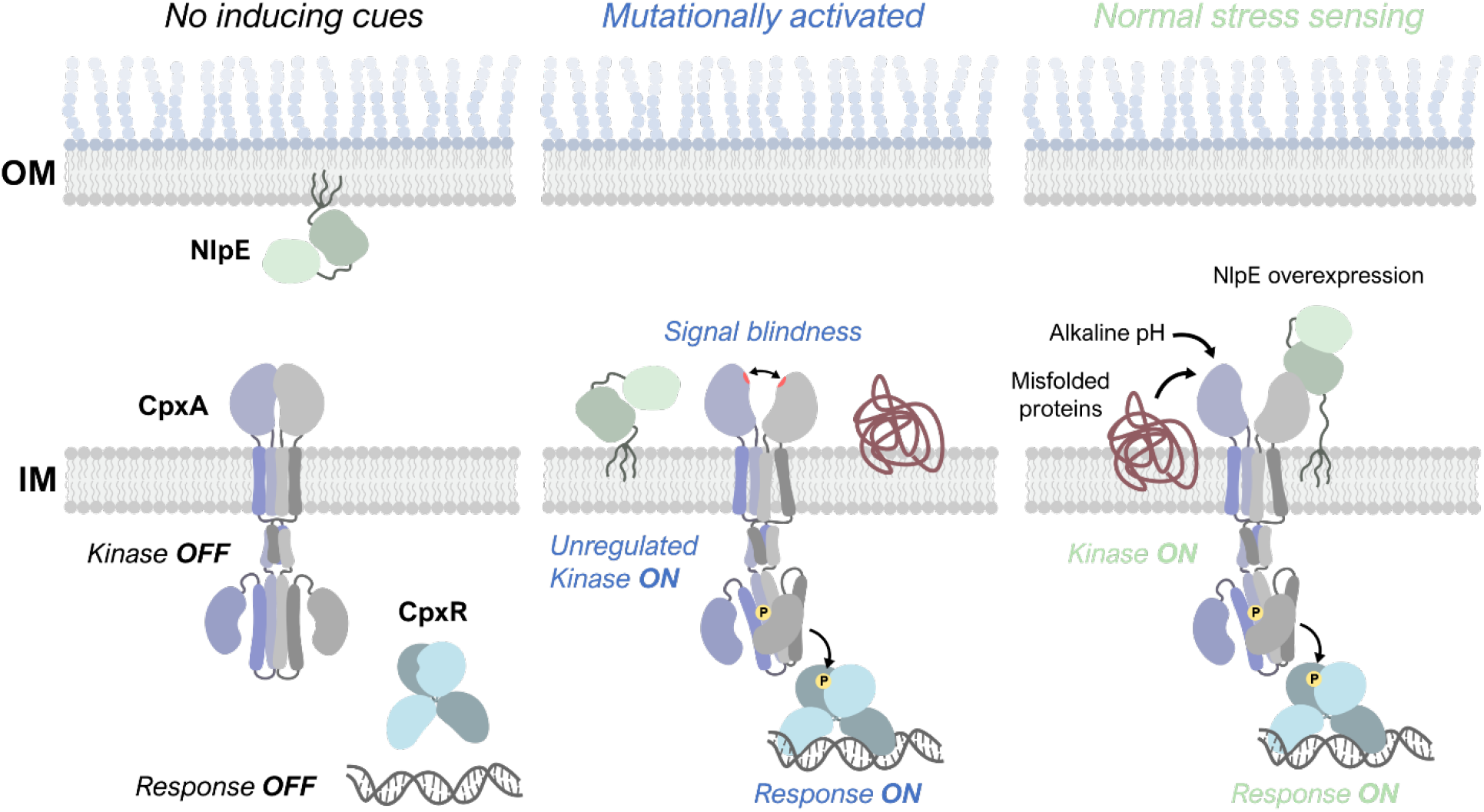
The novel organization of PAS domain dimers in CpxA regulates its activity. In the absence of inducing cues, CpxA is kept off by interactions between its PAS domains. Aberrant activation by mutations disrupting monomer-monomer interactions alter the structural relationship between the PAS and transmembrane domains. Thus, normal sensing CpxA likely involves relief of this auto-inhibition and transduction of signals through the PAS-TM interface.

Structural studies of histidine kinases have tended to focus on domains in isolation because of the modular organization of these proteins. The AlphaFold2 model of CpxA homodimers provides a basis to explain the phenotypes of previously described and novel mutations using a model of CpxA-SD in the context of its transmembrane and cytoplasmic domains. While experimentally determined, full-length structures of sensor kinases remain elusive, studies incorporating machine-learning based protein structure prediction tools and experimental approaches may provide a more holistic picture of how these proteins function and shed light on both the unity and diversity present in the broadly distributed family of PAS sensory folds.

## Experimental procedures

### Strains and growth conditions

All strains used in this study are listed in Table S2. Strains were cultured in lysogeny broth (LB; 10 g/L Bacto-Tryptone, 5 g/L Bacto-Yeast Extract, 5 g/L sodium chloride) with appropriate antibiotics (50 µg/ml kanamycin; 100 µg/ml ampicillin). Experiments involved in alkaline pH sensing used LB media containing 100 mM 3-(N-morpholino)propanesulfonic acid (MOPS) with pH adjusted with sodium hydroxide to pH 7.0 or 8.0 or LB media buffered with 100 mM sodium phosphate with pH adjusted to 5.8 (non-inducing) or 8.0 (inducing). A final concentration of 0.1 mM IPTG was used in experiments to induce NlpE or CpxA expression from plasmids. All cultures were grown at 37°C with shaking at 225 RPM.

### Strain construction

A list of primers used in this study are listed in Table S3. RM53 (TR50 Δ*cpxA*) was created by P1 generalized transduction using lysates generated from the Keio collection of mutants as previously described^70^. Kanamycin resistance cassettes were removed using plasmid-based expression of the Flp recombinase as previously described^71^. Strains encoding for *cpxA* chromosomal mutants N107A, K121A, and Y123A were generated by using site-directed mutagenesis and λRed recombination^72^. The native *cpxA* chromosomal locus was replaced with a counter-selectable *cat-sacB* cassette^73^, which was amplified with 50 bp of homology to the 5’ and 3’ ends of *cpxA* using the primers in Table S3, by an initial round of λRed recombination. Successful recombination was confirmed by chloramphenicol resistance and sucrose sensitivity.

A plasmid encoding *cpxA* with each of the aforementioned mutations were generated by replacing the *cpxA* locus of plasmid pCA-*cpxA* from the ASKA library in the pCA24N plasmid backbone^74^. Site-directed mutagenesis was conducted as described by Ko and Ma^75^. Briefly, the regions upstream and downstream of the codon to be mutated were amplified separately by PCR. Each product was amplified with one primer containing an SfiI site for cloning into pCA24N and the other being a mutagenic primer containing an EarI site (Table S3). Both products were digested with EarI and ligated to yield an SfiI-flanked product with the desired alanine substitution. This product was purified with a QIAquick PCR purification kit (QIAGEN), digested with SfiI, and ligated into SfiI-digested pCA-*cpxA* to yield pCA24N-based vectors with *cpxA* harbouring the desired mutations.

These plasmids were then used as templates to generate PCR fragments containing each *cpxA* variant flanked by about 40 bp to the 5’ and 3’ ends of the *cpxA*::*cat-sacB* region for a second round of λRed recombination. These fragments were then electroporated into MC4100 *cpxA*::*cat-sacB* containing the λRed system, and successful recombinants were selected by screening for growth on sucrose and sensitivity to chloramphenicol, and mutations were confirmed by sequencing. Finally, a *cpxP-lacZ* reporter carried on a recombinant λ phage was moved into all mutant strain backgrounds using two P1 transductions. First, a *nadA*::Tn10 allele tightly linked to the λ attachment site which confers resistance to tetracycline and the inability to grow on minimal media (MM) was transduced into the mutant strains. Tet^R^ and MM-transductants were used in a second transduction with P1 lysate prepared from TR50 (lRS88 *cpxP-lacZ*)^22^. Lac^+^ transductants were selected on minimal media containing X-gal, and these transductants were screened for sensitivity to tetracycline to yield the strains used in experiments.

### Plasmid construction

A list of plasmids used in this study are listed in Table S4. Expression vectors for CpxA were created through standard molecular biology techniques. For the crystallization construct, the predicted periplasmic domain of CpxA (amino acids 31-163) was amplified from *E. coli* MC4100 using primers 5’-31 and 3’-163 (Table S3) with BamHI and EcoRI restriction sites. Amplified *cpxA*_31-163_ was cloned into BamHI/EcoRI-digested pGEX-6P-1 expression vector (GE Healthcare), which encodes for an N-terminal glutathione S-transferase (GST) tag and PreScission^TM^ Protease cleavage sites.

The pK184-*cpxA* vectors used in reporter experiments was generated using restriction digest cloning procedures. Because *cpxA* contains an internal EcoRI cut site, plasmid pK184 was digested with EcoRI, treated with the Klenow fragment and religated to eliminate this site. *cpxA* coding sequence was amplified from pCA-*cpxA*^74^ with flanking BamHI-HindIII sites added using primers listed in Table S3. After PCR amplification, BamHI and HindIII (Invitrogen) digested *cpxA* PCR product was ligated into BamHI and HindIII digested pK184. *cpxA* ligation was confirmed by restriction digest analysis and Sanger sequencing (Molecular Biology Services Unit, University of Alberta).

Mutations were introduced into pK184-*cpxA* (except M48K and D113K, see below) by site-directed mutagenesis using the Q5 site-directed mutagenesis kit (New England Biolabs) according to manufacturer instructions. Mutagenesis primers were generated using NEBaseChanger (New England Biolabs) and PCR conditions were determined using recommended annealing temperatures. Plasmids pK184-*cpxA*_M48K_ and pK184-*cpxA*_D113K_ were generated by overlap extension PCR using standard molecular techniques using the primers listed in Table S3. All mutations were confirmed by Sanger sequencing (Molecular Biology Services Unit, University of Alberta).

### CpxA expression and purification

Plasmid pGEX-*cpxA* was transformed into *E. coli* BL21(DE3). Cells were cultured at 30°C in LB with ampicillin until the culture reached an optical density of 0.8-0.9 at 600 nm. Cultures were then induced with 0.05 mM IPTG and grown at 22°C for 21 hours. CpxA_31-163_-GST was purified at 4°C using a protocol modified from the GST Gene Fusion System Handbook (GE Healthcare). Briefly, cells were harvested by centrifugation, and pellets were resuspended in buffer (50 mM Tris-HCl pH 7.5, 400 mM NaCl, and protease inhibitors leupeptin, pepstatin, and phenylmethylsulfonyl fluoride). Cells were lysed using chicken egg white lysozyme treatment and sonication. Lysates were centrifuged and clarified; clarified lysate was mixed with Glutathione Sepharose 4B resin (GE Healthcare) which was pre-equilibrated in resuspension buffer. The binding reaction was incubated with nutation for 1.5 hrs. The resin was then washed with 10 volumes of resuspension buffer, and the bound CpxA-GST was eluted using resuspension buffer containing 20 mM reduced glutathione at pH 7.5.

The elutions were then pooled, buffer exchanged into resuspension buffer, and concentrated using a 10,000 MWCO centrifugal filter device (Millipore, Fisher Scientific). The GST tag was cleaved by PreScission Protease (GE Healthcare) treatment over 12 hrs at 4°C with the progress of reaction monitored by SDS-PAGE. The cleavage reaction was then applied to a second and third affinity column to remove resilient proteolyzed GST and any remaining fusion proteins. The column flow-through and washes containing CpxA_31-163_ were pooled and concentrated using a 3,000 MWCO spin concentrator (Millipore, Fisher Scientific). As a final purification step, CpxA_31-163_ was passed over a HiLoad Superdex 75 26/60 size-exclusion column equilibrated with resuspension buffer on an ÄKTA purifier (GE Healthcare). The CpxA_31-163_ elution peak fractions were pooled and concentrated using a 3,000 MWCO spin concentrator.

### Crystallization and data collection

Purified CpxA_31-163_ was dialyzed into crystallization buffer (50 mM HEPES pH 7.0, 350 mM NaCl) using 3000 MWCO spin concentrator. The concentration was estimated to be 21.1 mg/ml based on absorbance at 280 nm, using the theoretical extinction coefficient of 26,470 M^-1^cm^-1^ derived from the amino acid sequence of CpxA_31-163_ plus extra residues using ProtParam^76^. Initial crystallization conditions for CpxA_31-163_ were identified from the NeXtal JCSG+ ProComplex Suite, F3 (Qiagen) and were further optimized. CpxA_31-163_ was crystallized by hanging drop vapour diffusion at room temperature (∼21°C) by mixing 1 µl of protein at 21.1 mg/ml in crystallization buffer with 1 µl of reservoir solution (100 mM Tris-HCl pH 7.5, 20% v/v 2-methyl-2,4-pentanediol).

Native protein crystals use for phasing were formed in similar conditions by mixing 1 µl of CpxA at 21.1 mg/ml in crystallization buffer with 1 µl of reservoir solution. The crystals were harvested and flash frozen in liquid nitrogen without any additional cryoprotectant. Diffraction data were collected at the Advanced Light Source, SIBYLS beamline 12.3.1 (Berkeley, CA) and Canadian Light Source, beamline CMCF-BM 08b1-1 (Saskatoon, Canada). CpxA crystallized in the space group P212121 and diffraction data data was collected to a resolution of 1.8 Å.

### Structure solution and refinement

The periplasmic domain of *V. parahaemolyticus* CpxA (PDB 3V67)^23^ was used as a search model for molecular replacement. A single domain (Chain A, residues 45-159) was mutated to the corresponding *E. coli* sequence based on sequence alignment and further modified using Sculptor^77^ in PHENIX^78^ by pruning sidechains using the Schwarzenbacher method^79^. Molecular replacement with PHASER^80^ using data to 2.0 Å placed two copies with a TFZ sore of 8.0. A model of the same domain was generated using SWISSMODEL^81,82^, superimposed on the molecular replacement solution and improved with iterative model building in PHENIX AutoBuild^83^ followed by refinement in phenix.refine^84^ resulting in an R-free of 0.323. One of the monomers was then used as the search model to phase the 1.8 Å data placing two copies with a PHASER TFZ sore of 52.1. Subsequent iterative manual model building in COOT^85^ and refinement in PHENIX yielded a final model with R-free 0.247.

### Generation of multiple sequence alignments (MSA) of *cpxA* for AlphaFold2 modelling

The query sequence, residues 1-184 of *E. coli* CpxA (Uniprot P0AE82), was input into pHMMER (https://www.ebi.ac.uk/Tools/hmmer/search/phmmer) and searched against the uniprotKB database with default search settings, except BLOSUM45 was used, yielding an MSA of 2379 sequences. Jalview was used to remove all sequences with 98% or 96% redundancy resulting in MSAs of 580 and 321 sequences, respectively. The 98% redundant MSA was made gapless for the *E. coli* CpxA sequence and used for AlphaFold2 modeling (pHMMER_98%_non-redundant). The 96% redundant MSA was realigned in Jalview using the MAFFT E-INS-i preset, since the middle (PAS) sequence was known to have low sequence homology^10^. This realigned MSA was used for all analysis.

### AlphaFold2 modeling of CpxA-SD

CpxA structural modeling was performed using the ColabFold implementation of AlphaFold2^43,44^. Sequences of either *E. coli* or *V. parahaemolyticus* CpxA were input into the AlphaFold2_MMseqs2 Google Colab notebook (v 1.3.0), and multiple sequence alignments were generated using only the MMseqs2 algorithm or were supplemented with a pHMMER MSA (see MSA generation). All runs used the mulitmer_V2 weights, 48 recycles and generated five models; full input parameters are listed in Table S5. The models were ranked by pTM score, and the highest scoring model was used for analysis. Pymol (Version 2.4.0, Schrödinger, LLC) was used for analyzing the AlphaFold2 models and for creating images. Confidence metrics were analyzed using python3 scripts and plotted with Gnuplot (Version 5.4, http://www.gnuplot.info/).

### β-galactosidase assays

Activation of the Cpx response was measured by quantifying the activity of a chromosomal *cpxP-lacZ* reporter. Strains prepared in biological triplicate were subcultured into LB with kanamycin (and ampicillin for NlpE overexpression experiments) from overnight cultures for 2 hours. For alkaline pH experiments, cells were pelleted by centrifugation at 4000 RPM for 10 minutes and resuspended in LB+100mM MOPS pH 7.0 or 8.0 and grown for a further 2 hours. For NlpE overexpression experiments, cells were induced with 0.1 mM IPTG and grown for a further 3 hours with shaking. β-galactosidase assays were conducted as previously described^34,86^. Briefly, cultures were centrifuged at 4000 RPM for 10 minutes, and pellets were resuspended in Z-buffer^87^. Optical density (absorbance at 600nm) was measured from an aliquot of resuspended culture for standardization. Each culture was treated with chloroform and sodium dodecyl sulfate (SDS) and vortexed to permeabilize cells. Absorbance at 420nm (A_420_) was measured in a plate reader for each culture at 30 second intervals after the addition of 10 mg/ml ONPG. β-galactosidase activity (*cpxP-lacZ* reporter) activity was quantified by calculating the maximum slope of the A_420_ measures standardized to that of the respective culture’s optical density as previously described. Statistical significance was using unpaired *t-* tests (Prism, GraphPad).

### SDS-PAGE and Western blotting

SDS-PAGE and Western blotting was used to confirm expression of CpxA from our plasmids according to standard procedures. Cultures were grown in the same conditions as for other experiments. The optical density of each culture was measured and the equivalent of OD 4.0 cells in 100 µl was collected. Cells were washed twice in phosphate buffered saline (PBS) and resuspended in water and 2× Laemmli buffer (Sigma). Samples were heated at 95°C for 5 minutes and cooled to room temperature before being separated on an 8% tris-glycine SDS-PAGE gel and transferred to a nitrocellulose membrane using a semi-dry transfer machine (Bio-Rad). Membranes were blocked in 5% skim milk dissolved in Tris-buffered saline with 0.1% Tween-20 (TBST) for 1-3 hours and incubated with anti-CpxA-MBP antibody^22^ in 2% bovine serum albumin (BSA) in TBST overnight. The next day, membranes were washed four times with TBST and incubated with fluorescent IRDye680RD (goat anti-mouse) antibodies in 5% milk in TBST for 1 hour and imaged using a Bio-Rad ChemiDoc imager.

## Data Availability

All data found in this manuscript can be shared upon request to T.L.R. The structure of CpxA-SD_EC_ have been deposited to the RCSB PDB (PDB ID 8UK7). All code used in this manuscript have been deposited temporarily at links found in the “Supporting Information” document and a permanent DOI will be made available after acceptance.

This article contains supporting information which can be found in the “Supporting Information” document.

## Acknowledgements and Funding Information

Diffraction data were collected at the Advanced Light Source, SIBYLS beamline 12.3.1 (Berkeley, CA) and Canadian Light Source, beamline CMCF-BM 08b1-1 (Saskatoon, Canada). This work was supported by operating grants from the Natural Sciences and Engineering Research Council (NSERC RGPIN-2021-02710 to T.L.R. and NSERC Discovery Grant RGPIN-2016-05163 to J.N.M.G.) and the Canadian Institutes of Health Research (CIHR MOP 142347 to T.L.R. and CIHR168972 to J.N.M.G.). T.H.S.C. is supported by the NSERC Postgraduate Scholarship (PGS-D) award. G.L.T. is supported by an AHFMR Studentship Award and a recipient of the CIHR Frederick Banting and Charles Best CGS Doctoral Award.

## CRediT Author Statement

**Timothy H. S. Cho:** Conceptualization, Investigation, Visualization, Methodology, Writing – Original Draft **Cameron Murray:** Conceptualization, Investigation, Software, Methodology, Writing – Original Draft **Roxana Malpica:** Methodology, Investigation **Rodrigo Margain-Quevedo:** Methodology, Investigation **Gina L. Thede:** Methodology, Investigation **Jun Lu:** Investigation **Ross A. Edwards:** Methodology, Formal analysis **J. N. Mark Glover:** Supervision, Writing – Review & Editing, Funding acquisition **Tracy L. Raivio:** Supervision, Writing – Review & Editing, Funding acquisition

