## Supplemental information for "The sensor of the bacterial histidine kinase CpxA is a novel dimer of extracytoplasmic Per-ARNT-Sim (PAS) domains"

**Table S1.** Data collection and refinement statistics

| | CpxA (31-163) | CpxA (31-163) † |
|---|---|---|
| **Wavelength** | 1.03316 | 0.97950 |
| **Resolution range** | 39.2-1.8 (1.86-1.8) | 39.6-2.0 (2.07-2.0) |
| **Space group** | P 21 21 21 | P 21 21 21 |
| **Unit cell** | 37.16 43.16 186.98 90 90 90 | 37.05 43.77 186.34 90 90 90 |
| **Total reflections** | 227024 (21825) | 170877 (5615) |
| **Unique reflections** | 28482 (2769) | 21058 (1864) |
| **Multiplicity** | 8.0 (7.9) | 8.1 (3.0) |
| **Completeness (%)** | 98.1 (96.6) | 98.4 (89.0) |
| **Mean I/sigma(I)** | 13.75 (2.68) | 13.83 (1.37) |
| **Wilson B-factor** | 34.85 | 34.28 |
| **R-merge** | 0.070 (1.09) | 0.078 (0.800) |
| **R-meas** | 0.076 (1.16) | 0.083 (0.926) |
| **R-pim** | 0.028 (0.408) | 0.027 (0.488) |
| **CC 1/2** | 0.997 (0.958) | 0.999 (0.741) |
| **CC*** | 0.999 (0.989) | 1 (0.923) |
| **Reflections used in refinement** | 28337 (2735) | 21045 (1861) |
| **Reflections used for R-free** | 1345 (133) | 969 (86) |
| **R-work** | 0.204 (0.303) | |
| **R-free** | 0.247 (0.348) | |
| **CC(work)** | 0.955 (0.919) | |
| **CC(free)** | 0.954 (0.856) | |
| **Number of non-hydrogen atoms** | 2032 | |
| macromolecules | 1943 | |
| solvent | 89 | |
| **Protein residues** | 233 | |
| **RMS(bonds)** | 0.013 | |
| **RMS(angles)** | 1.15 | |
| **Ramachandran favored (%)** | 97.82 | |
| **Ramachandran allowed (%)** | 2.18 | |
| **Ramachandran outliers (%)** | 0.00 | |
| **Rotamer outliers (%)** | 2.91 | |
| **Clashscore** | 3.10 | |
| **Average B-factor** | 56.92 | |
| macromolecules | 57.16 | |
| solvent | 51.77 | |
| **Number of TLS groups** | 16 | |

Statistics for the highest-resolution shell are shown in parentheses.

† Used for phasing.

**Table S2.** Strains used in this study.

| Strain | Description | Source |
|---|---|---|
| MC4100 | F_ *araD139* (*argF-lac*)*U169 rpsL150* (Strr) *relA1 flbB5301 decC1 ptsF25 rbsR* | (1) |
| TR50 | MC4100 λRS88[*cpxP'-lacZ*$^{+}$] | (2) |
| RM53 | TR50 Δ*cpxA* | This study |
| GLT100 | BL21(DE3) + pGEX-*cpxA*$_{31\text{-}163}$ | This study |
| RM336 | MC4100 *cpxA*$_{N107A}$ | This study |
| RM367 | MC4100 *cpxA*$_{K121A}$ | This study |
| RM338 | MC4100 *cpxA*$_{Y123A}$ | This study |
| RM441 | MC4100 IRS88[*cpxP-lacZ*] *cpxA*$_{N107A}$ | This study |
| RM448 | MC4100 IRS88[*cpxP-lacZ*] *cpxA*$_{K121A}$ | This study |
| RM444 | MC4100 IRS88[*cpxP-lacZ*] *cpxA*$_{Y123A}$ | This study |
| RM477 | RM441 + pBR322 | This study |
| RM478 | RM441 + pLD404 | This study |
| RM481 | RM448 + pBR322 | This study |
| RM482 | RM448 + pLD404 | This study |
| RM483 | RM444 + pBR322 | This study |
| RM484 | RM444 + pLD404 | This study |
| TC636 | RM53 + pK184 | This study |
| RMQ2 | RM53 + pK184-*cpxA*$_{WT}$ | This study |
| RMQ6 | RM53 + pK184-*cpxA*$_{M48K}$ | This study |
| RMQ7 | RM53 + pK184-*cpxA*$_{D113K}$ | This study |
| RMQ21 | RM53 + pK184-*cpxA*$_{WT}$ + pCA24N | This study |
| RMQ22 | RM53 + pK184-*cpxA*$_{WT}$ + pCA-*nlpE* | This study |
| RMQ23 | RM53 + pK184-*traJ* + pCA24N | This study |
| RMQ24 | RM53 + pK184-*traJ* + pCA-*nlpE* | This study |
| RMQ27 | RM53 + pK184-*cpxA*$_{M48K}$ + pCA24N | This study |
| RMQ28 | RM53 + pK184-*cpxA*$_{M48K}$ + pCA-*nlpE* | This study |
| RMQ29 | RM53 + pK184-*cpxA*$_{D113K}$ + pCA24N | This study |
| RMQ30 | RM53 + pK184-*cpxA*$_{D113K}$ + pCA-*nlpE* | This study |
| RMQ34 | RM53 + pK184-*cpxA*$_{E91A}$ | This study |
| RMQ52 | RM53 + pK184-*cpxA*$_{E91A}$ + pCA24N | This study |
| RMQ53 | RM53 + pK184-*cpxA*$_{E91A}$ + pCA-*nlpE* | This study |
| TC643 | RM53 + pK184-*cpxA*$_{E91K}$ | This study |
| TC644 | RM53 + pK184-*cpxA*$_{E91K+R93E}$ | This study |
| TC646 | RM53 + pK184 + pTrc99A | This study |
| TC647 | RM53 + pK184 + pTrc-*nlpE* | This study |
| TC648 | RM53 + pK184-*cpxA* + pTrc99A | This study |
| TC649 | RM53 + pK184-*cpxA* + pTrc-*nlpE* | This study |
| TC650 | RM53 + pK184-*cpxA*$_{E91K}$ + pTrc99A | This study |
| TC651 | RM53 + pK184-*cpxA*$_{E91K}$ + pTrc-*nlpE* | This study |
| TC652 | RM53 + pK184-*cpxA*$_{E91K+R93E}$ + pTrc99A | This study |
| TC653 | RM53 + pK184-*cpxA*$_{E91K+R93E}$ + pTrc-*nlpE* | This study |
| TC726 | RM53 + pK184-*cpxA*$_{N107D}$ | This study |
| TC719 | RM53 + pK184-*cpxA*$_{Q103E}$ | This study |
| TC721 | RM53 + pK184-*cpxA*$_{R106E}$ | This study |
| TC758 | RM53 + pK184-*cpxA*$_{N107D}$ + pTrc99A | This study |
| TC761 | RM53 + pK184-*cpxA*$_{N107D}$ + pTrc-*nlpE* | This study |
| TC763 | RM53 + pK184-*cpxA*$_{Q103E+D113N}$ | This study |
| TC756 | RM53 + pK184-*cpxA*$_{Q103E}$ + pTrc99A | This study |
| TC759 | RM53 + pK184-*cpxA*$_{Q103E}$ + pTrc-*nlpE* | This study |
| TC757 | RM53 + pK184-*cpxA*$_{R106E}$ + pTrc99A | This study |
| TC760 | RM53 + pK184-*cpxA*$_{R106E}$ + pTrc-*nlpE* | This study |

| | | |
|---|---|---|
| TC724 | RM53 + pK184-*cpxA*$_{E91K+R99E}$ | This study |
| TC725 | RM53 + pK184-*cpxA*$_{E91K+R93E+R99E}$ | This study |
| TC796 | RM53 + pK184-*cpxA*$_{E91K+R99E}$ + pTrc99A | This study |
| TC797 | RM53 + pK184-*cpxA*$_{E91K+R99E}$ + pTrc-*nlpE* | This study |
| TC798 | RM53 + pK184-*cpxA*$_{E91K+R93E+R99E}$ + pTrc99A | This study |
| TC799 | RM53 + pK184-*cpxA*$_{E91K+R93E+R99E}$ + pTrc-*nlpE* | This study |

**Table S3.** Primers used in this study.

| Primer Name | Sequence (5'-3') | Notes |
|---|---|---|
| *For generation of* cpxA *chromosomal mutants* | | |
| pCAF8 | CGTCTTCACCTCGAGAAATC | Anchor primer to pCA24N encoding SfiI sites |
| pCAR4 | TTGCATCACCTTCACCCTCTCCACTGACAG | Anchor primer to pCA24N encoding SfiI sites |
| CpxAN$_{107}$AFw | AACTCTTCAGCTTTTATTGGTCAGGCCGA | Mutagenic primer to generate N107A fragment with EarI site |
| CpxAN$_{107}$ARv | AACTCTTCAAGCACGAATGATCTGCATTTCG | Mutagenic primer to generate N107A fragment with EarI site |
| CpxAK$_{121}$AFw | AACTCTTCAGCTAAGTATGGCCGCGTGGA | Mutagenic primer to generate K121A fragment with EarI site |
| CpxAK$_{121}$ARv | AACTCTTCAAGCCTTCTGCGGATGATCGG | Mutagenic primer to generate K121A fragment with EarI site |
| CpxAY$_{123}$AFw | AACTCTTCAGCAGGCCGCGTGGAACTGGT | Mutagenic primer to generate Y123A fragment with EarI site |
| CpxAY$_{123}$ARv | AACTCTTCATGCCTTTTTCTTCTGCGGAT | Mutagenic primer to generate Y123A fragment with EarI site |
| *cpxA-cat* | ATTTAATGTGGTGGCGGCGTCTGTTCCGGGCGATTG ATAAGTGGGCACCGTGTGACGGAAGATCACTTCGCAG | For amplification of *cat-sacB* cassette |
| *sacB-cpxA* | GGTCAAACAGTAAGTTAATGAAATCGGATTGAGAA CTGCTGGCCGGATCAAAGGGAAAACTGTCCATAT | For amplification of *cat-sacB* cassette |
| *cpxA*181Fw | TCCGCCCAACGATTTAATG | For amplification of *cpxA* variants from pCA24N expression vector |
| *cpxA*503Rv | AGCAGTAATAGCGGGCGGT | For amplification of *cpxA* variants from pCA24N expression vector |
| *For generation of CpxA$_{31-163}$ expression vector* | | |
| 5'-31 | CGCGGATCCGATTCACGCCAGATGACCGA | For amplification of *cpxA*$_{31-163}$ from genome |
| 3'-163 | GCGAATTCCTAGCGGTCAAACAGTAAGTT | For amplification of *cpxA*$_{31-163}$ from genome |
| *For generation of pK184*-cpxA | | |
| XA-1 fw | ATAGGATCCGTGAGGAGGTTCCTATGATAGGCAGCTTAACCGCGC | For amplification of *cpxA* with EcoRI site (and for creation of overlap fragments, see below) |
| XA-2c rv | TATATAAGCTTCTGCAGTTATGACCGCTTATACAGCGGCAACCAAATCACC | For amplification of *cpxA* with BamHI site |
| pK184_F | CGTATGTTGTGTGGAATTGTG | For sequencing of inserts into pK184 |
| pK184_R | CAAGGCGATTAAGTTGGGTAA | For sequencing of inserts into pK184 |
| *For generation of mutations in pK184*-cpxA | | |
| XA-2b rv | GGCAAGGAATTCCTGTGGCCC | For generation of downstream fragment for all mutations |
| XA-5 fw | AGATTGAGCAGCATGTCGAAGCG | For generation of downstream fragment (with XA-2b) to create M48K mutation by overlap extension PCR |
| XA-6 rv | CGCTTCGACATGCTGCTCAATCTTCAGACCCTGACGCTGTTCGCT | For generation of upstream fragment (with XA-1) to create M48K mutation by overlap extension PCR |

| | | |
|---|---|---|
| XA-7 fw | AAAAACGCCGATCATCCGCAGAAG | For generation of downstream fragment (with XA-2b) to create D113K mutation by overlap extension PCR |
| XA-8 rv | CTTCTGCGATGATCGGCGTTTTTGGCCTGACCAATAAAGTTACGAATGATCTGC | For generation of upstream fragment (with XA-1) to create D113K mutation by overlap extension PCR |
| CpxA E91A-fw | GGTGACCACCGCTGGCCGCGTGA | For generation of E91A by Q5 site-directed mutagenesis |
| CpxA E91A-rv | AATAACAAACGCTGTCCTGGC | For generation of E91A by Q5 site-directed mutagenesis |
| CpxA E91K-fw | GGTGACCACCAAAGGCCGCGTGA | For generation of E91K by Q5 site-directed mutagenesis |
| CpxA E91K-rv | AATAACAAACGCTGTCCTGGCGG | For generation of E91K by Q5 site-directed mutagenesis |
| Q5SDM_E91K_R93E_F | CGAAGTGATCGGCGCTGAACGC | For generation of E91K+R93E by Q5 site-directed mutagenesis |
| Q5SDM_E91K_R93E_R | CCTTTGGTGGTCACCAATAACAAACGC | For generation of E91K+R93E by Q5 site-directed mutagenesis |
| Q5SDM_99+91_F | CGGCGCTGAAGAAAGCGAAATGCAGATCATTC | For generation of E91K+R99E by Q5 site-directed mutagenesis |
| Q5SDM_99+91_R | ATCACGCGGCCTTTGGTG | For generation of E91K+R99E by Q5 site-directed mutagenesis |
| Q5SDM_91 93 99_F | CGGCGCTGAAGAAAGCGAAATGC | For generation of E91K+R93E+R99E by Q5 site-directed mutagenesis |
| Q5SDM_91 93 99_R | ATCACTTCGCCTTTGGTG | For generation of E91K+R93E+R99E by Q5 site-directed mutagenesis |
| Q5SDM_Q103E_F | CAGCGAAATGGAAATCATTCGTAACTTTATTG | For generation of Q103E by Q5 site-directed mutagenesis |
| Q5SDM_Q103E_R | CGTTCAGCGCCGATCACG | For generation of Q103E by Q5 site-directed mutagenesis |
| Q5SDM_R106E_F | GCAGATCATTGAAAACTTTATTGGTCAGGCC | For generation of R106E by Q5 site-directed mutagenesis |
| Q5SDM_R106E_R | ATTTCGCTGCGTTCAGCG | For generation of R106E by Q5 site-directed mutagenesis |
| Q5SDM_N107D_F | GATCATTCGTGATTTTATTGGTCAGGCC | For generation of N107D by Q5 site-directed mutagenesis |
| Q5SDM_N107D_R | TGCATTTCGCTGCGTTCA | For generation of N107D by Q5 site-directed mutagenesis |
| Q5SDM_D113N_F | TGGTCAGGCCAATAACGCCGATC | For generation of Q103E+D113N by Q5 site-directed mutagenesis |
| Q5SDM_103+113_R | ATAAAGTTACGAATGATTTCCATTTCGC | For generation of Q103E+D113N by Q5 site-directed mutagenesis |

**Table S4.** Plasmids used in this study.

| Plasmid | Description | Source |
|---|---|---|
| pCA24N | Empty ASKA library vector, $Cam^R$ | |
| pCA-*cpxA* | CpxA expression from pCA24N backbone, IPTG-inducible, ASKA library (GFP-), $Cam^R$ | |
| pTrc99A | Empty expression vector, IPTG-inducible from *trc* promoter, $Amp^R$ | |
| pTrc-$nlpE_{WT}$ | His-tagged NlpE expression from pTrc99A backbone, IPTG-inducible, $Amp^R$ | |
| pBR322 | Cloning vector, $Amp^R$ | |
| pLD404 | NlpE expression from the pBR322 backbone, $Amp^R$ | |
| pK184 | Empty expression vector, $Kan^R$ | |
| pK184-*traJ* | pK184 encoding for the *traJ* locus which was used as a less toxic vector control in some experiments as it has no impact on activation of the Cpx response, $Kan^R$ | |
| pK184-$cpxA_{WT}$ | WT *cpxA* cloned into pK184, $Kan^R$ | This study |
| pK184-$cpxA_{M48K}$ | *cpxA* M48K cloned into pK184, $Kan^R$ | This study |
| pK184-$cpxA_{D113K}$ | *cpxA* D113K cloned into pK184, $Kan^R$ | This study |
| pK184-$cpxA_{E91A}$ | *cpxA* E91A cloned into pK184, $Kan^R$ | This study |
| pK184-$cpxA_{E91K}$ | *cpxA* E91K cloned into pK184, $Kan^R$ | This study |
| pK184-$cpxA_{E91K+R93E}$ | *cpxA* E91K+R93E cloned into pK184, $Kan^R$ | This study |
| pK184-$cpxA_{N107D}$ | *cpxA* N107D cloned into pK184, $Kan^R$ | This study |
| pK184-$cpxA_{Q103E}$ | *cpxA* Q103E cloned into pK184, $Kan^R$ | This study |
| pK184-$cpxA_{R106E}$ | *cpxA* R106E cloned into pK184, $Kan^R$ | This study |
| pK184-$cpxA_{Q103E+D113N}$ | *cpxA* Q103E+D113N cloned into pK184, $Kan^R$ | This study |
| pK184-$cpxA_{E91K+R99E}$ | *cpxA* E91K+R99E cloned into pK184, $Kan^R$ | This study |
| pK184-$cpxA_{E91K+R93E+R99E}$ | *cpxA* E91K+R93E+R99E cloned into pK184, $Kan^R$ | This study |
| pFLP2 | Plasmid encoding for Flp recombinase, $Amp^R$ | |
| pKD46 | Plasmid encoding for λRed functions, $Amp^R$ | |
| pRM24 | pCA-$cpxA_{N107A}$, pCA24N vector harbouring *cpxA* N107A | This study |
| pRM12 | pCA-$cpxA_{K121A}$, pCA24N vector harbouring *cpxA* K121A | This study |
| pRM13 | pCA-$cpxA_{Y123A}$, pCA24N vector harbouring *cpxA* Y123A | This study |

**Table S5.** Modeling parameters and outputs of ColabFold.

| Model | Avg pLDDT | pTm | Sequences used | # Sequences | Start Res | End Res |
|---|---|---|---|---|---|---|
| CpxA ecoli Dimer_10 | 87.65 | 0.76 | mmSeqs2 + pHMMER_98%_non-redundant | 1214 | 8 | 184 |
| CpxA vib | 87.09 | 0.82 | mmSeqs2 | 484 | 14 | 195 |
| CpxA ecoli PAS-TM-HAMP | 85.69 | 0.76 | mmSeqs2 | 3647 | 1 | 233 |

A

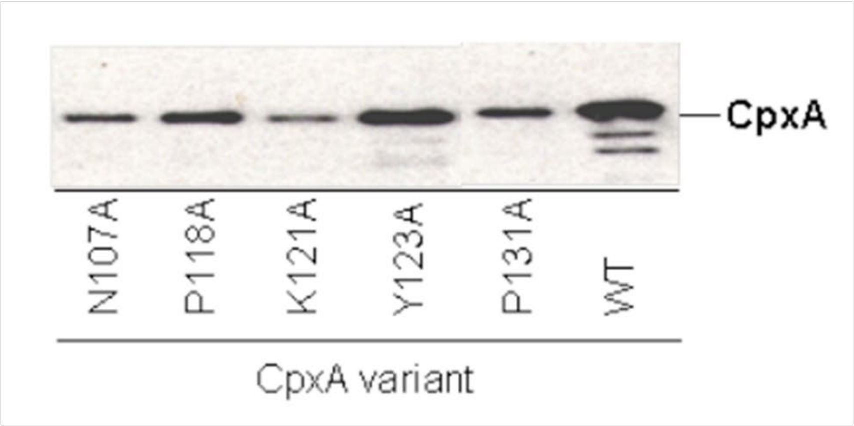


B

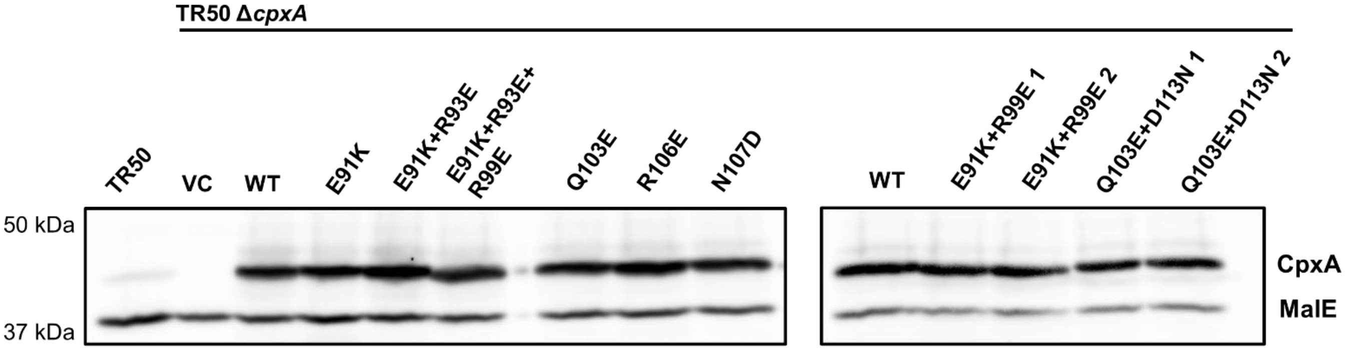


**Figure S1.** Expression levels of **(A)** chromosomal mutants and **(B)** plasmid-based mutations of CpxA. For chromosomal mutants, whole membrane fractions were purified and blotted for expression levels of CpxA using an anti-CpxA-MBP antibody. For plasmid-based CpxA, whole cell lysates were used for Western blotting.

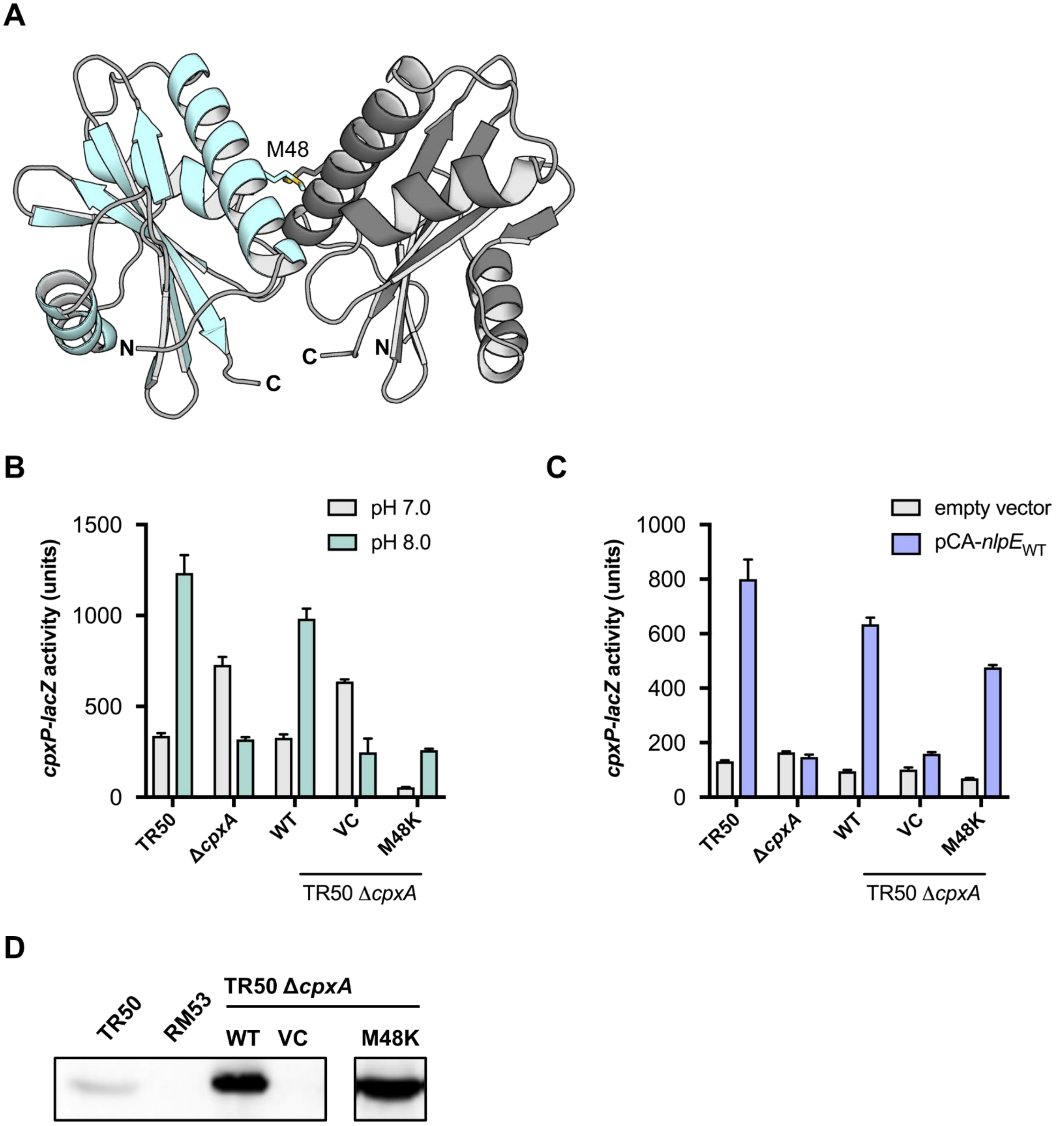


**Figure S2.** Crystal dimer structure of CpxA. **(A)** Ribbon cartoon diagram of the dimer with each monomer shown in a different color. The main dimer interface residue M48 is highlighted. The ability of the M48K mutation to sense **(B)** alkaline pH and **(C)** NlpE overexpression, as seen in the activity of a *cpxP-lacZ* transcriptional reporter. **(D)** shows the expression levels of CpxA in relevant strains as determined by Western blotting with anti-CpxA-MBP antibody.

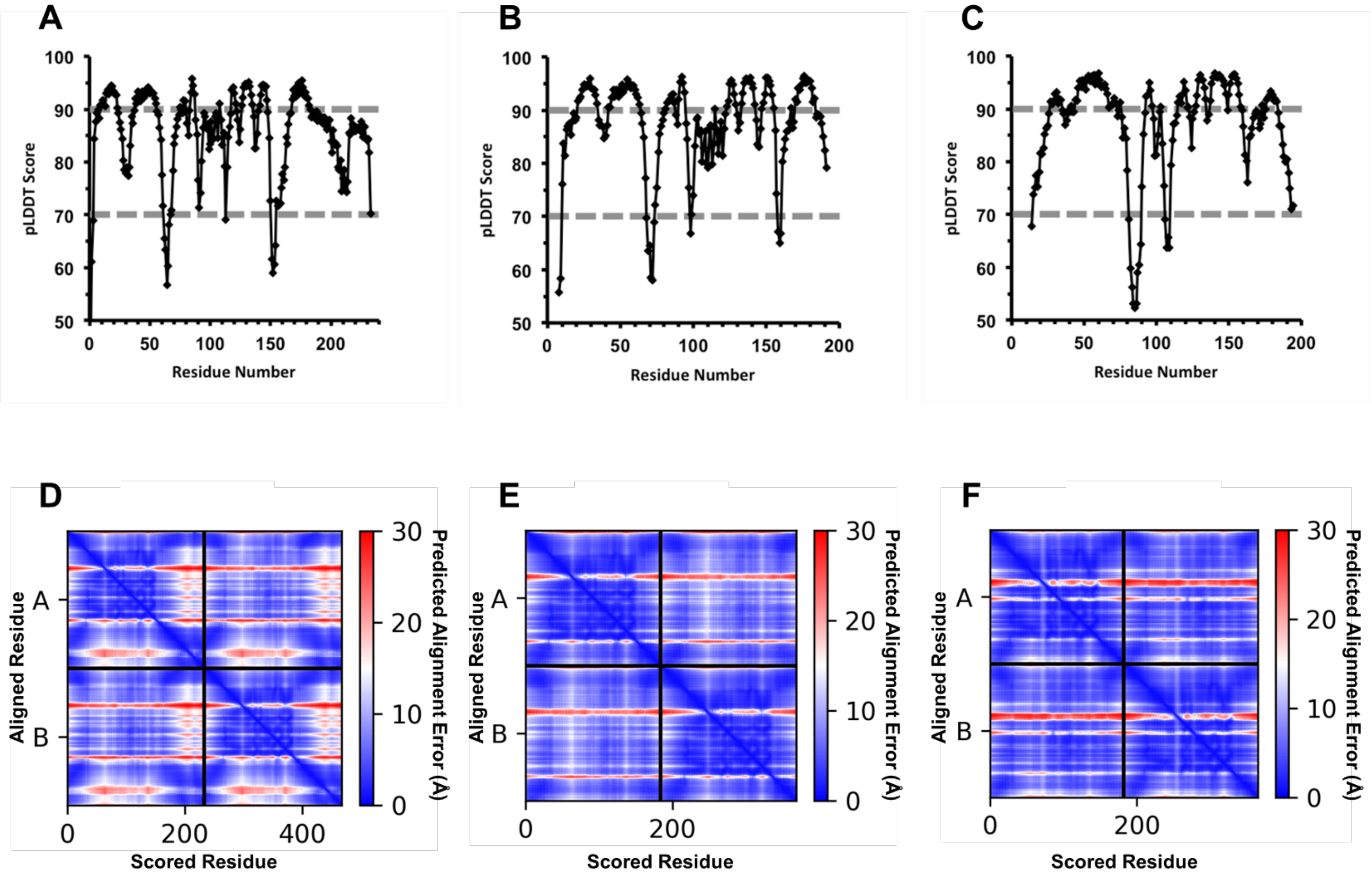


**Figure S3.** AlphaFold2 confidence metrics for *E. coli* and *V. parahaemolyticus* models. **A-C** Predicted local distance difference test (pLDDT) scores per residue for CpxA-EC TM1-HAMP **(A)**, CpxA-SD$_{EC}$ **(B)**, CpxA-SD$_{Vib}$ **(C)**. Dashed lines indicate cut offs for very high confidence (pLDDT > 90) and high confidence (pLDDT > 70). **D-F** Predicted alignment error (pAE) within monomers (Top Left and Bottom Right sub-panels) and between monomers (Top Right and Bottom Left sub-panels) for CpxA-EC TM1-HAMP **(D)**, CpxA-SD$_{EC}$ **(E)**, CpxA-SD$_{Vib}$ **(F)**.

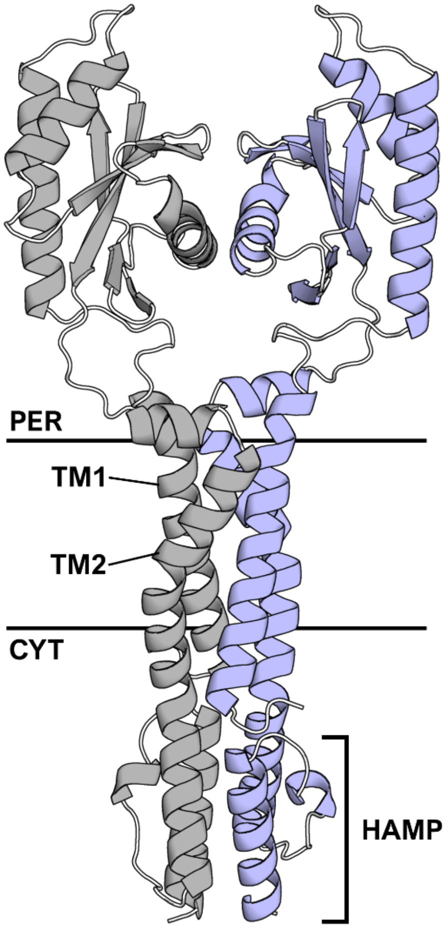


**Figure S4.** AlphaFold2 model of *E. coli* CpxA including transmembrane domains, periplasmic sensor domains and cytosolic HAMP domains.

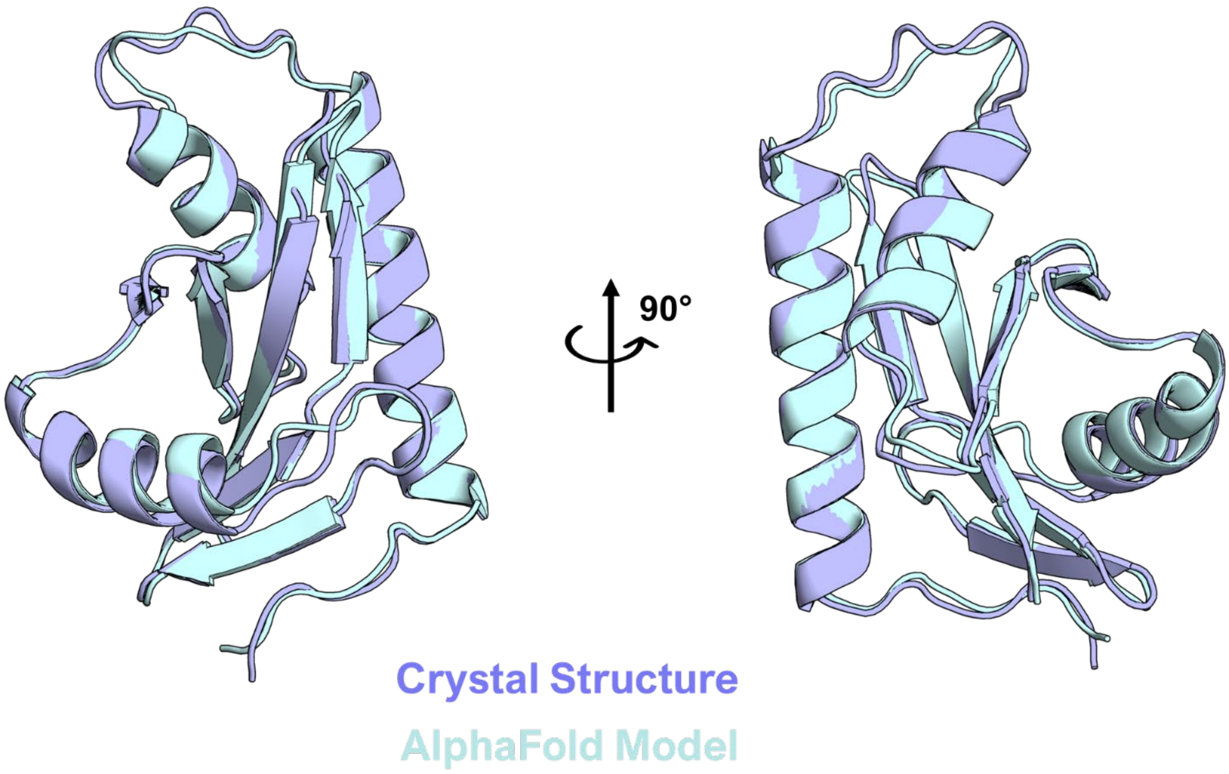


**Figure S5.** Alignment of the crystal structure and AlphaFold2 model monomer of *E. coli* $CpxA_{SD}$.

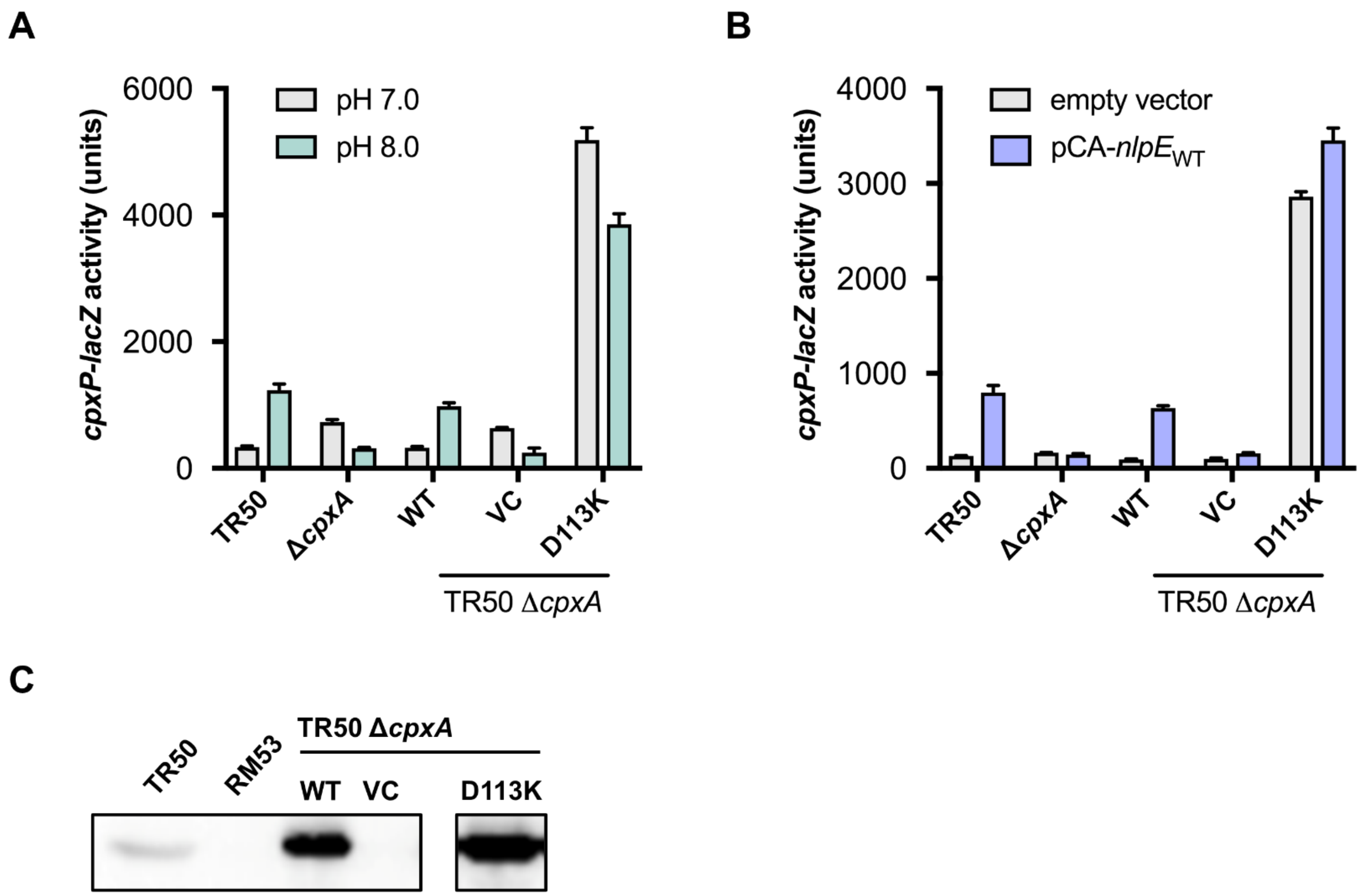


**Figure S6.** Ability of plasmid-borne CpxA D113K variant to sense **(A)** alkaline pH and **(B)** NlpE overexpression. **(C)** shows the expression level of D113K compared to WT CpxA by Western blotting.

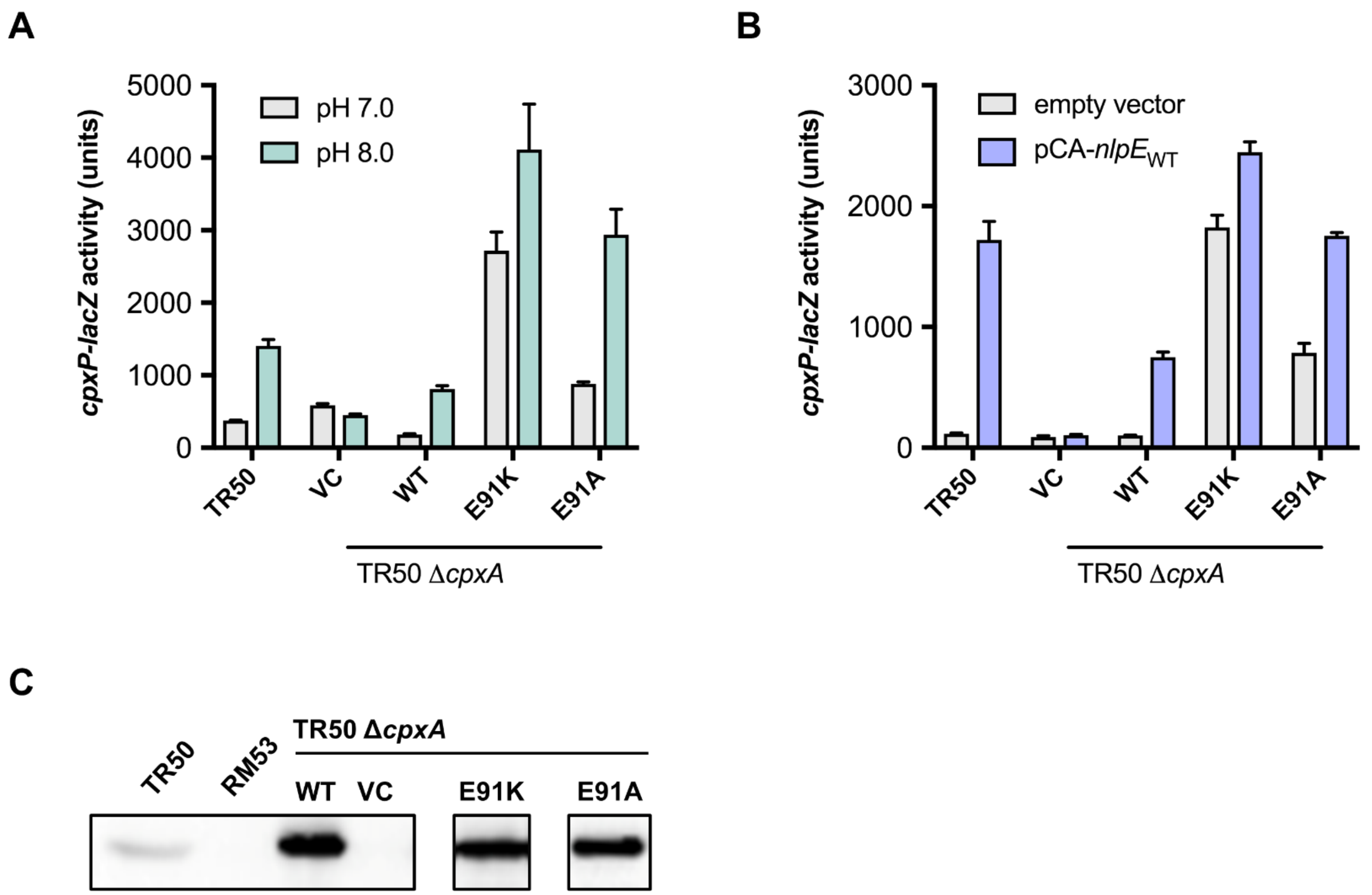


**Figure S7.** Ability of plasmid-borne CpxA E91K and E91A variants to sense **(A)** alkaline pH and **(B)** NlpE overexpression. **(C)** shows the expression level of D113K compared to WT CpxA by Western blotting.

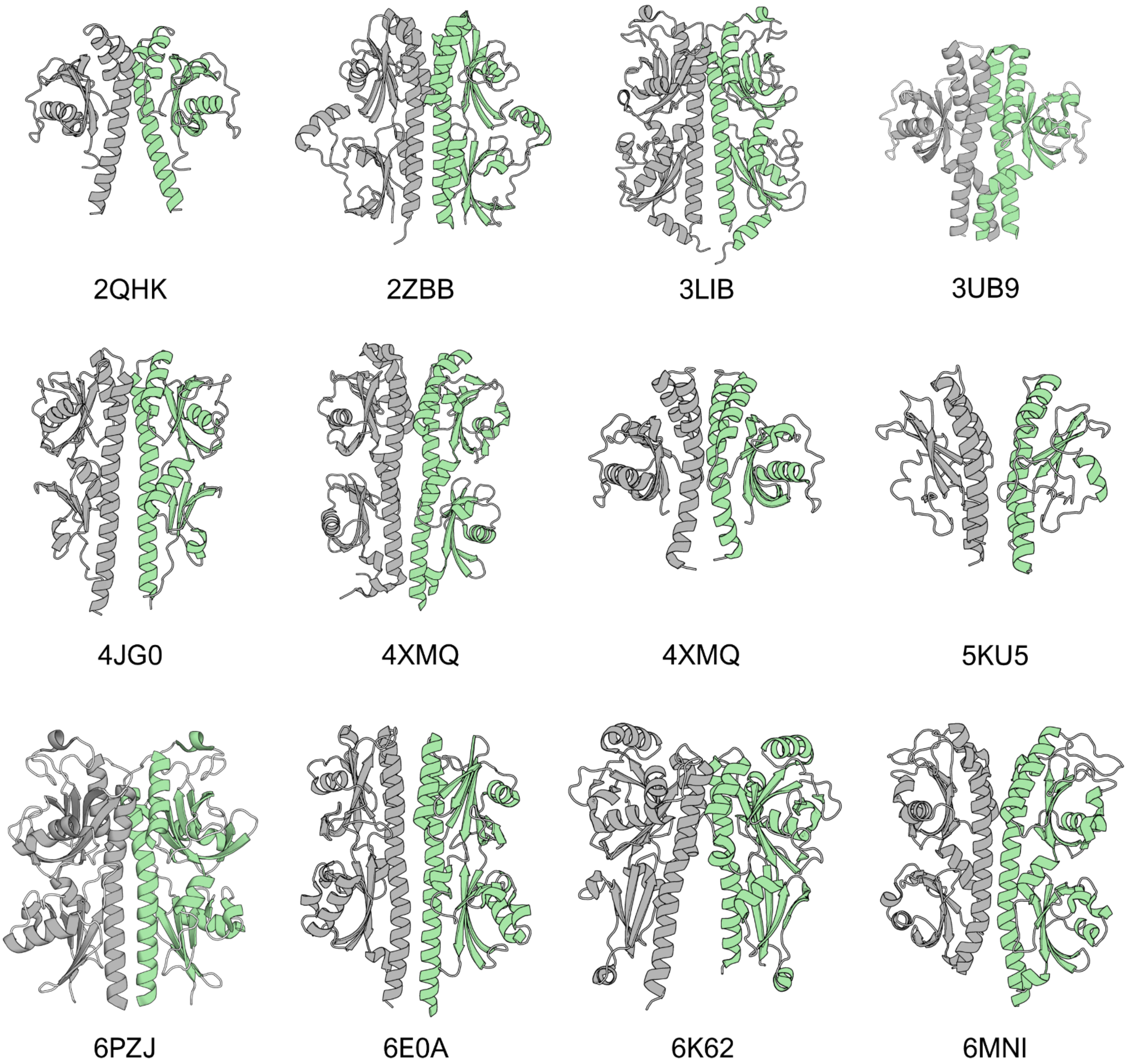


**Figure S8.** More PAS domain dimer hits of $CpxA_{SD}$. Each monomer is represented as a different colored chain (grey vs green). Protein Database (PDB) codes for each structure are listed below.

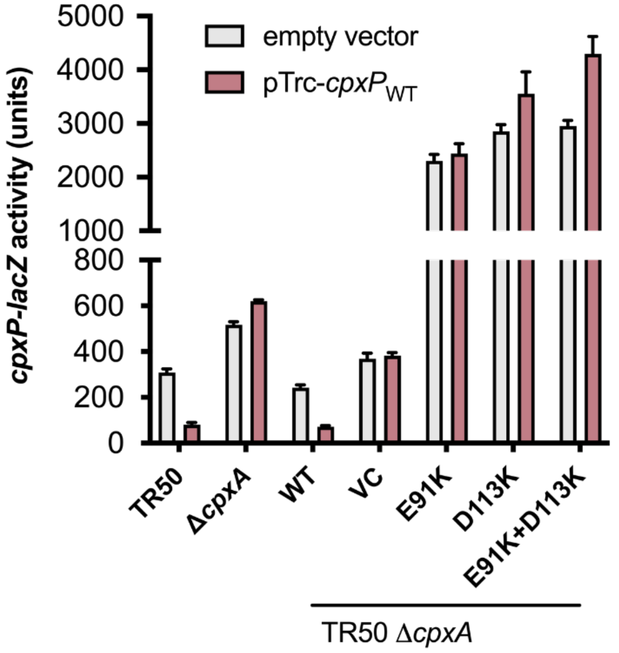


**Figure S9.** The ability of hyperactivated CpxA variants to sense CpxP overexpression. CpxP was induced from plasmid pTrc-*cpxP* with 0.1 mM IPTG for 2 hours after cells reached mid-log phase. The activity of a *cpxP-lacZ* reporter was used to measure activation of CpxA. Indicated CpxA variants were expressed from plasmid pK184.

**A**

| | ...[EQH] | ...[DN] | Other | Total |
|---|---|---|---|---|
| P... | 93 | 6 | 56 | 155 |
| [STDN]... | 42 | 9 | 21 | 72 |
| [QE]... | 24 | 4 | 7 | 35 |
| Other | 45 | 14 | 0 | 59 |
| Total | 204 | 33 | 84 | 321 |

**B**

| | ...[EQH] | ...[DN] | Other | Total |
|---|---|---|---|---|
| P... | 0 | 0 | 0 | 0 |
| [STDN]... | 3 | 0 | 63 | 65 |
| [QE]... | 1 | 0 | 109 | 110 |
| Other | 16 | 2 | 128 | 146 |
| Total | 20 | 2 | 299 | 321 |

**Figure S10.** Conservation of N-capping motifs in *cpxA* sequences. **(A)** shows sequence motifs that are present at the N-cap site. **(B)** shows the lack of the presence of N-capping motifs in sequences immediately following the N-cap position in CpxA-$SD_{EC}$ and CpxA-$SD_{Vib}$.

This readme is for analyzing the dali results of CpxA.

Required Files:

Original Files:

parseDali.py (makes the hit dictionary) (https://pastebin.com/wmcR46k5)

makePymolSessions.py (performs search) (https://pastebin.com/UHC0xXzZ)

Internet Scripts:

https://pymolwiki.org/index.php/AngleBetweenHelices

http://pymolwiki.org/index.php/RotationAxis

Installation:

For ease of use all required files should be in the same directory. Files in other directories, such as the pymolwiki scripts, can be imported via the sys.path.insert line in makePymolSessions.py.

Execution:

This script requires pymol to be downloaded. Edit the input lines in the makePymolSessions.py file (searchPDB, readFile, writeDir) to the desired file paths and optionally modify any of the search parameters (a3HelixDistanceCutoff, etc.). Run the makePymolSessions.py file via

pymol -cr makePymolSession.py
